# Determinants in gammaretroviral Env dictate the production of neutralizing antibodies

**DOI:** 10.64898/2026.07.30.741819

**Authors:** Robert Z. Zhang, Lia Robben, Ashley Willey, Vincent Mele, Emilia Crow, Sandra Umana, Brady Callahan, Laura Arroyo, Grace W. Graudin, Holly C. Simmons, Kevin R. McCarthy, Melissa Kane

## Abstract

Protective immune responses are shaped by the nature of the infectious pathogen. Some viral infections are self-limiting and stimulate lasting immunity, while others activate poor responses that fail to control the infection or prevent development of disease. Little is known concerning the specific viral determinants that control whether a given viral infection will result in protective immunity. Here, we demonstrate that strains of the gammaretrovirus, murine leukemia virus, differ in their ability to stimulate neutralizing antibody production via a noncanonical pathway. Additionally, we show that virus-specific production of neutralizing immune responses is unique to infection, suggesting that the activation of alternative antibody responses is triggered by the infectious process rather than the unique nature of specific viral antigens. Viral chimeras indicate that activation of neutralizing antiviral antibody production is determined by the receptor-binding domain, and that the surface subunit controls binding to and infection of various cell types. The ability of specific viral factors to determine the outcome of infection and the induction of distinctive immune responses between closely related retroviral strains provides an ideal model for the dissection of both host and viral determinants underlying the control of viral infection and the production of neutralizing antibodies.

**SUMMARY:** We report here that retroviral envelope glycoprotein determines stimulation of GC B cell responses and the production of neutralizing antibodies in the absence of IFNγ-signaling, thus defining the factors that stimulate non-canonical immune responses and further developing the knowledge required for vaccine development.

## INTRODUCTION

Humoral immunity in the form of protective antibodies serves as a critical barrier to prevent viral pathogenesis (Burton, 2002; Corti and Lanzavecchia, 2013). Model organisms such as inbred mice have proved to be invaluable for investigation of the pathways underlying protective humoral immune responses against viruses due to the inability to perform genetic manipulations in humans. In this regard, murine retroviruses, such as the gammaretrovirus murine leukemia virus (MLV), have provided essential insights into the molecular mechanisms underlying antiviral immune responses. MLV is transmitted as an exogenous virus through bodily fluids or as endogenous stably integrated provirus and primarily infects cells of lymphoid origin (Portis et al., 1987; Rosenberg and Jolicoeur, 1997). MLVs compose a large family of retroviruses, and individual strain isolates often induce different disease phenotypes upon infection of susceptible strains of mice (Friend, 1957; Gross, 1951; Hook et al., 2002; Moloney, 1960; Rauscher, 1962; Rowe and Hartley, 1972).

An essential step in the development of effective humoral responses is cytokine-induced class switch recombination (CSR), which results in the production of specific antibody isotypes (Stavnezer et al., 2008). Importantly, the enrichment of specific isotypes following infection influences the effectiveness of antiviral immune responses due to functional differences determined by the constant region, including the capacity to activate complement (Klaus et al., 1979), phagocytosis-activating Fc receptors on macrophages (Heusser et al., 1977), and induction of antibody-dependent cellular cytotoxicity (ADCC) (Kipps et al., 1985). *In vivo*, CSR requires B cell receptor stimulation, Toll-like receptor signaling, CD40 signaling, as well as interactions with helper T cells and the activation of isotype-specific promoters by cytokines (Stavnezer and Schrader, 2014). CSR is mediated by transcription from isotype-specific promoters upstream of the C_H_ genes followed by recruitment of activation-induced cytidine deaminase (AID) that initiates an irreversible intrachromosomal recombination (Muramatsu et al., 2000; Muramatsu et al., 1999). Canonical antiviral antibody responses in mice involve an IFNγ-dependent CSR that results in the production of immunoglobulins of the IgG2a isotype (Snapper and Paul, 1987). However, some viruses and parasites have been shown to stimulate neutralizing IgG2a antibodies via a non-canonical IFNγ-independent pathway (Kane et al., 2018; Markine-Goriaynoff et al., 2000; McKendall and Woo, 1988; Nguyen et al., 1994; van den Broek et al., 1995; Zhang et al., 2023), but neither the mechanisms by which isotype-specific promoters are activated nor the additional cytokines that promote class switch recombination in the IFNγ-independent pathway are fully understood. While the direct comparison of human and mouse immunoglobulin subtypes is not possible, the capacity of cytokines to direct isotype selection and affect the outcome of infections is common to both mice and humans (Vidarsson et al., 2014). The dominant antiviral isotype in humans is IgG1 (Huang et al., 2006; Nachbagauer et al., 2016; Siemoneit et al., 1994), although the bias to this isotype is not as strong as the IgG2a bias in inbred mice. No human IgG1-specific “switch factor” has been described (Pan-Hammarstrom et al., 2007), and antiviral antibody production in humans involves a pathway distinct from the canonical IFNγ-dependent pathway observed in most inbred mice. Thus, the non-canonical IFNγ-independent pathway for the production of neutralizing antiviral antibodies in mice may be more reflective of the signaling networks that direct humoral immune responses in humans.

Using two closely related strains of MLV, Rauscher-like (RL) and Moloney (Mo), we found that RL-MLV infection, but not Mo-MLV infection, generates neutralizing antiviral antibodies in the absence of IFNγ signaling. Further, Mo- and RL-MLV infection stimulate distinct antibody isotype profiles. We report that protective IFNγ-independent antibody responses are controlled by determinants in the gammaretroviral envelope protein. We demonstrate that the alternative pathway for protective antibodies is specific to responses stimulated by infection and is determined by the receptor-binding domain (RBD) of gammaretroviral Env. Sequence differences in the surface-subunit (SU) determine binding affinity to various cell types; however, binding affinity only correlated with productive infection of Gr-1^+^ myeloid cells.

## Results

### IFNγ-independent antiviral antibody responses control RL-MLV, but not Mo-MLV infection

We have previously reported that RL-MLV infection stimulates neutralizing antibodies in both IFNγ-deficient BALB.J (BALB/cJ mice congenic for the *H2^j^* MHC haplotype) and IFNγR-deficient 129S7 mice (Kane et al., 2018; Zhang et al., 2023). To investigate whether the production of protective IFNγ-independent immunity is specific to RL-MLV infection, we infected IFNγ-deficient BALB.J mice and IFNγR-deficient 129S7 mice with Mo-MLV. Infected mice were then monitored for neutralizing antibody production and viral titers in the spleen. In contrast to RL-MLV infection, which stimulated the production of protective neutralizing antibodies in all animals **(Fig. 1A-C)**, mice infected with Mo-MLV produced significantly lower titers of anti-MLV antibodies in the absence of IFNγ signaling **(Fig. 1D)**. Antibodies produced against Mo-MLV by either IFNγ- or IFNγR-deficient mice were not neutralizing (high pAUC), and these animals failed to clear the infection **(Fig. 1E-F)**. Since the antibodies produced upon Mo-MLV infection in IFNγ-deficient mice were non-neutralizing, we investigated the antiviral antibody isotype profile of both Mo- and RL-MLV infected mice against the SU of the MLV envelope protein (Env) **(Fig. 1G)**. We found that IFNγ-deficient mice infected with Mo-MLV failed to efficiently generate IgG class-switched antibodies against MLV-SU. These results indicate that RL-MLV, but not Mo-MLV, infection stimulates class-switched antibodies via the noncanonical IFNγ-independent pathway. Since MLV SU is a critical target for neutralizing antibodies, we next sought to determine whether anti-SU antibodies are cross-neutralizing. Sera from WT mice infected with Mo-MLV recognized RL-SU **(Fig. 1H)** but did not neutralize RL-MLV **(Fig. 1I)**. Notably, both WT and IFNγ-deficient mice infected with RL-MLV generated anti-SU antibodies that exhibited reduced cross-reactivity against Mo-SU **(Fig. 1H)** and did not neutralize Mo-MLV **(Fig. 1I)**. These data demonstrate that SU-specific antibodies elicited upon MLV infection can recognize epitopes common between these MLV strains but fail to cross-neutralize and that strains of MLV differ in their ability to stimulate IFNγ-independent antibody production.

**Figure 1.**
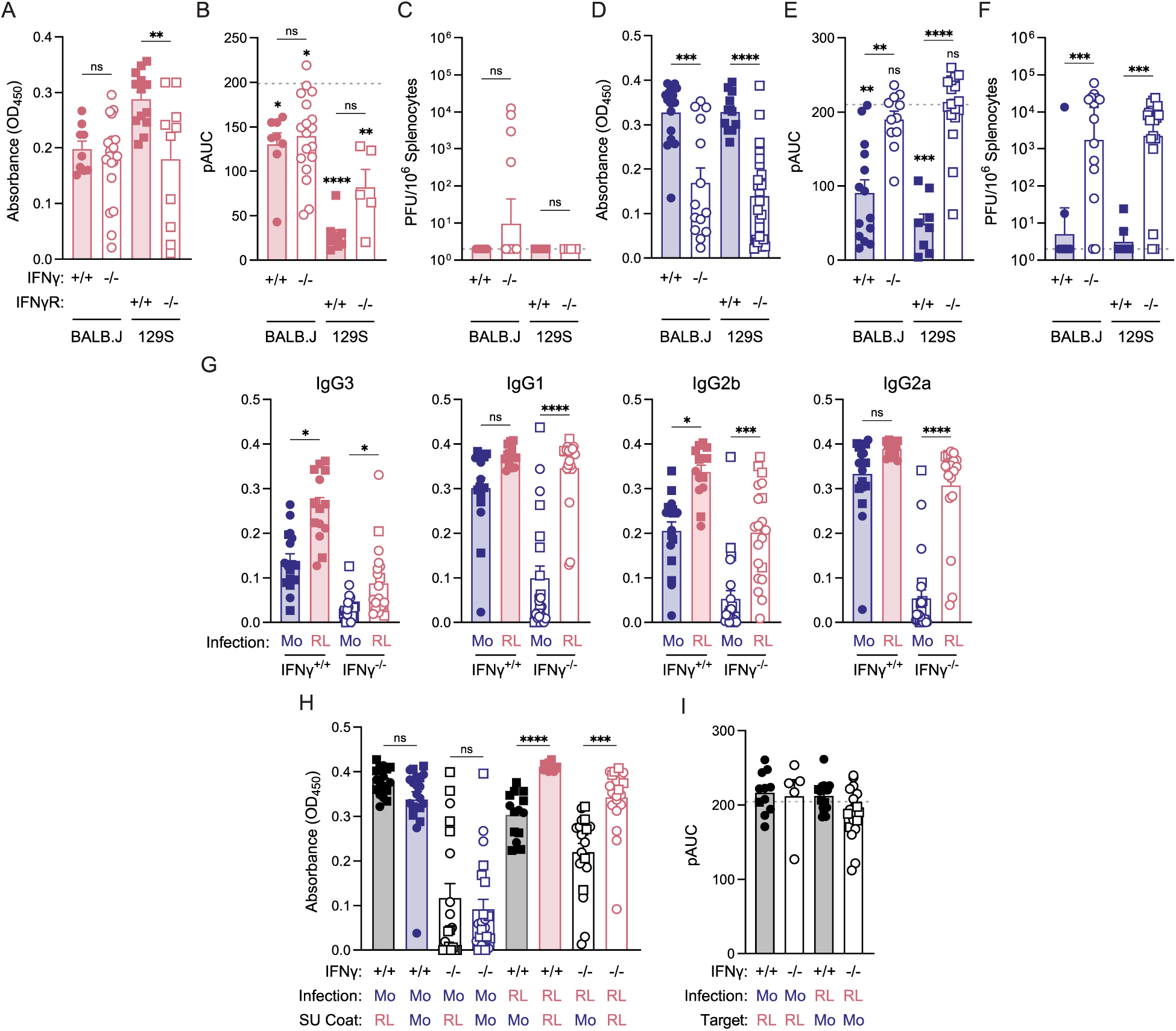
IFNγ-independent antiviral antibody responses control RL-MLV, but not Mo-MLV infection. Mice of the indicated genotypes were infected with RL-MLV **(A-C)** or Mo-MLV **(D-F)** and analyzed eight weeks post infection. **(A)** Sera was collected and monitored for total Igs against RL-MLV virion proteins. **(B)** Sera was serially diluted and incubated with RL-MLV before adding to SC-1 cells. Neutralization capacity was quantified by determining the partial area under the curve (pAUC). Grey dashed line indicates the average pAUC of naïve sera. Lower pAUC indicates more neutralization. Statistics above columns represent comparisons against naïve sera. **(C)** Splenocytes were isolated and subjected to an infectious center assay. Grey dashed line represents the limit of detection. **(D)** Antibody titers were measured as in (A) against Mo-MLV virion proteins. **(E)** Neutralization of Mo-MLV was measured as in (B). **(F)** Virus titers measured as in (C). **(G)** Sera from infected mice were monitored for IgG3, IgG1, IgG2b, and IgG2a antibodies against Mo- or RL-SU by ELISA. **(H)** Sera from infected mice were monitored for total Igs against the indicated MLV-SU. **(I)** Neutralization of the indicated viral infections against target viruses as measured in (B). Grey dashed line indicates the average pAUC of naïve sera. *For each statistical comparison, a nonparametric Kruskal-Wallis test was applied, and corresponding significance values are indicated for each graph. ns, not significant; *, p<0.05; **, p<0.01; ***, p<0.001; ****, p<0.0001*.

### RL-MLV and Mo-MLV infection stimulate distinct immune responses

Given that RL-MLV, but not Mo-MLV, infection stimulated neutralizing antibodies in the absence of IFNγ, we hypothesized that Mo-MLV infection may result in attenuated IFNγ-independent B cell responses. As there is no exact WT control for IFNγR-deficient 129S7 mice and to eliminate the influence of genetic differences between the 129S and BALB.J strains of mice, we focused on BALB.J mice for subsequent experiments. To analyze the B cell response upon MLV infection, we assessed germinal center (GC) responses in the spleen over five weeks by flow cytometry **(Fig. S1)**. In WT BALB.J mice, both Mo- and RL-MLV infection induced peak GC B cell responses at four weeks post infection **(Fig. S2A)**. Similar to WT mice, IFNγ-deficient mice infected with RL-MLV produced peak GC B cells at four weeks; however, IFNγ-deficient mice infected with Mo-MLV did not exhibit a significant expansion of GC B cells at any time point **(Figs. 2A** and **S2B)**. As the GC response peaked at four weeks post infection, we utilized this timepoint for further investigations of the antiviral response. We found that in both WT and IFNγ-deficient mice, Mo- and RL-MLV infection resulted in distinct isotype profiles in GC B cells **(Fig. 2B)**. Specifically, RL-MLV infection in WT mice induced a higher frequency of IgG1^+^ GC B cells compared to Mo-MLV infection. Additionally, in IFNγ-deficient mice, Mo-MLV infection preferentially induced IgG2b class switching while RL-MLV infection induced IgG2a class switching. While total plasmablast (PB) numbers were equivalent between the groups **(Fig. 2C)**, the frequency of IgG2a^+^ PBs was higher in RL-MLV infected mice **(Fig. 2D)**. Since GC reactions are dependent on CD4^+^ T helper cell interactions (Allen et al., 1993; Jacobson et al., 1974), we next investigated antiviral T cell responses. While we observed no difference in the total number of CD4^+^ T cells **(Fig. 2E)** or T follicular helper (Tfh) cells **(Fig. 2F)** in mice infected with Mo- or RL-MLV, we found that in most IFNγ-deficient mice, Mo-MLV infection induced more T follicular regulatory (Tfr) cells **(Fig. 2G)** but not total regulatory T cells (Tregs) compared to RL-MLV infection **(Fig. 2H)**. Accordingly, IFNγ-deficient RL-MLV infected mice had significantly higher ratios of Tfh:Tfr when compared to Mo-MLV infected mice **(Fig. 2I)**. These results suggest that, in the absence of IFNγ signaling, Mo-MLV infection results in a more immunosuppressive GC environment, thus impairing GC B cell expansion. Furthermore, in both IFNγ-sufficient and -deficient mice, T helper 1 (Th1) cell numbers were significantly higher upon Mo-MLV infection compared to RL-MLV infection **(Fig. 2J)**, suggesting that Mo-MLV infection is stimulating Th1 responses and that these Th1 responses are not dependent on IFNγ signaling.

**Figure 2.**
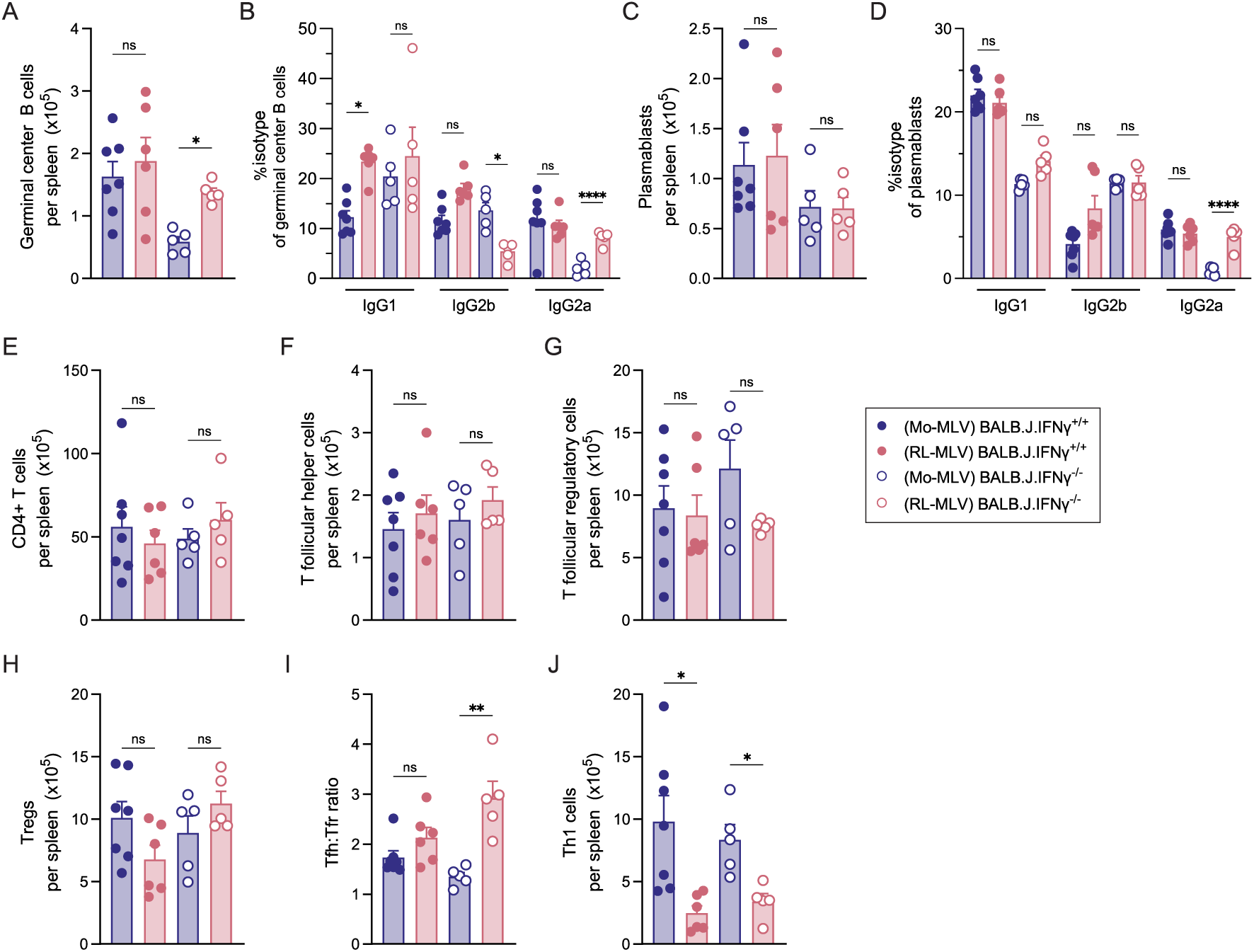
Mo- and RL-MLV infection stimulate distinct antiviral immune responses. Mice of the indicated genotypes were infected with either Mo-MLV or RL-MLV and splenocytes were isolated and analyzed by flow cytometry four weeks post infection. **(A)** Number of germinal center (GC) B cells. **(B)** Frequency of GC B cells that were IgM^−^IgG1^+^, IgM^−^IgG2b^+^, and IgM^−^ IgG2a^+^. **(C)** Number of plasmablasts (PB). **(D)** Frequency of PB that were IgM^−^IgG1^+^, IgM^−^IgG2b^+^, and IgM^−^IgG2a^+^. **(E)** Number of CD4+ T cells. **(F)** Number of T follicular regulatory (Tfr) cells. **(G)** Number of T follicular helper (Tfh) cells. **(H)** Ratio of Tfh to Tfr cells. **(I)** Number of T regulatory (Tregs) cells. **(J)** Number of T helper 1 (Th1) cells. *For each statistical comparison, a nonparametric Kruskal-Wallis test was applied, and corresponding significance values are indicated for each graph. ns, not significant; *, p<0.05; **, p<0.01; ****, p<0.0001*.

### IFNγ-independent immune responses are specifically stimulated by infection

To rule out the possibility that differences in the inherent antigenicity of Mo- and RL-MLV viral proteins control the induction of antiviral immune responses via the non-canonical IFNγ-independent pathway, we immunized both IFNγ-sufficient and -deficient mice with inactivated MLV virions in Complete Freund’s Adjuvant (CFA) and measured B and T cell responses at 12 days post immunization. We observed no difference in serum antibody isotypes IgM, IgG1, IgG3, IgG2b, or IgG2a against MLV virion proteins in mice immunized with Mo- or RL-MLV **(Fig. 3A)**. Additionally, we found no difference in GC B cells **(Fig. 3B-C)** or PBs **(Fig. 3D-E)** in mice immunized with RL- or Mo-MLV. IFNγ-deficient mice immunized with Mo-MLV had higher counts of CD4^+^ T cells **(Fig. 3F)** and Tregs **(Fig. 3G)**; however, there were no differences in the number of Tfr **(Fig. 3H)**, Tfh **(Fig. 3I)**, or Th1 cells **(Fig. 3J)** between the two groups. Thus, these results demonstrate that the distinct antiviral B cell and Tfh/Tfr responses against RL- and Mo-MLV are not mediated by antigenic differences between the two viral strains.

**Figure 3.**
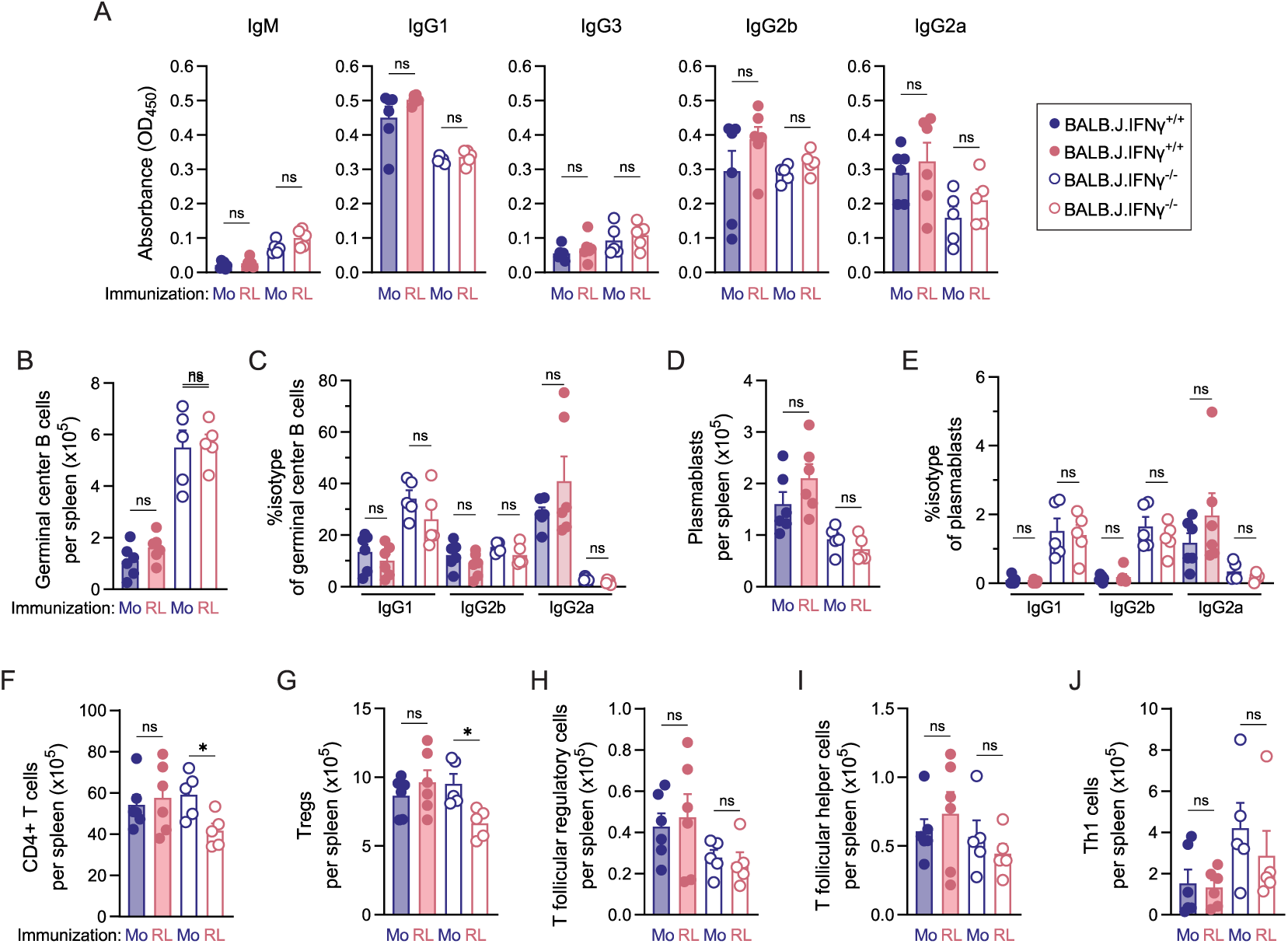
Differences in antiviral immune responses are specific to infection. Mice of the indicated genotypes were immunized with Triton X-100 treated Mo-MLV or RL-MLV and analyzed twelve days post immunization. **(A)** Sera was collected and monitored for IgM, IgG1, IgG3, IgG2b, and IgG2a antibodies against respective SU protein. **(B-J)** Splenocytes were isolated and analyzed by flow cytometry. **(B)** Number of germinal center (GC) B cells. **(C)** Frequency of GC B cells that were IgM^−^IgG1^+^, IgM^−^IgG2b^+^, and IgM^−^IgG2a^+^. **(D)** Number of plasmablasts (PB). **(E)** Frequency of PB that were IgM^−^IgG1^+^, IgM^−^IgG2b^+^, and IgM^−^IgG2a^+^. **(F)** Number of CD4+ T cells. **(G)** Number of T regulatory (Tregs) cells. **(H)** Number of T follicular regulatory (Tfr) cells. **(I)** Number of T follicular helper (Tfh) cells. **(J)** Number of T helper 1 (Th1) cells. *For each statistical comparison, a nonparametric Kruskal-Wallis test was applied, and corresponding significance values are indicated for each graph. ns, not significant; *, p<0.05*.

To determine whether differences in activating the IFNγ-independent pathway are due to a stimulatory signal unique to RL-MLV infection, or a suppressive signal induced by Mo-MLV infection, we coinfected IFNγ-deficient and -sufficient BALB.J mice with Mo- and RL-MLV and monitored them for neutralizing antibody production and viral load in the spleen. Coinfected mice cleared the infection **(Fig. 4A)**, generated neutralizing antibodies against both Mo- and RL-MLV **(Fig. 4B)**, although antibody titers against Mo-SU were reduced compared to RL-SU **(Fig. 4C-D),** suggesting that RL-MLV infection stimulates the IFNγ-independent production of neutralizing antibodies. Similar to singular infection with RL-MLV in IFNγ-deficient mice, coinfected mice generated GC responses **(Fig. 4E)** and had high ratios of Tfh cells to Tfr cells **(Fig. 4F)**. While coinfection induced Th1 responses, the responses were more similar to RL-MLV infection than Mo-MLV infection **(Fig. 4G)**, suggesting that the elevated Th1 responses observed during Mo-MLV infection were not generated in the presence of RL-MLV infection. Together, these data indicate that RL-MLV infection actively stimulates the IFNγ-independent neutralizing antibody response pathway.

**Figure 4.**
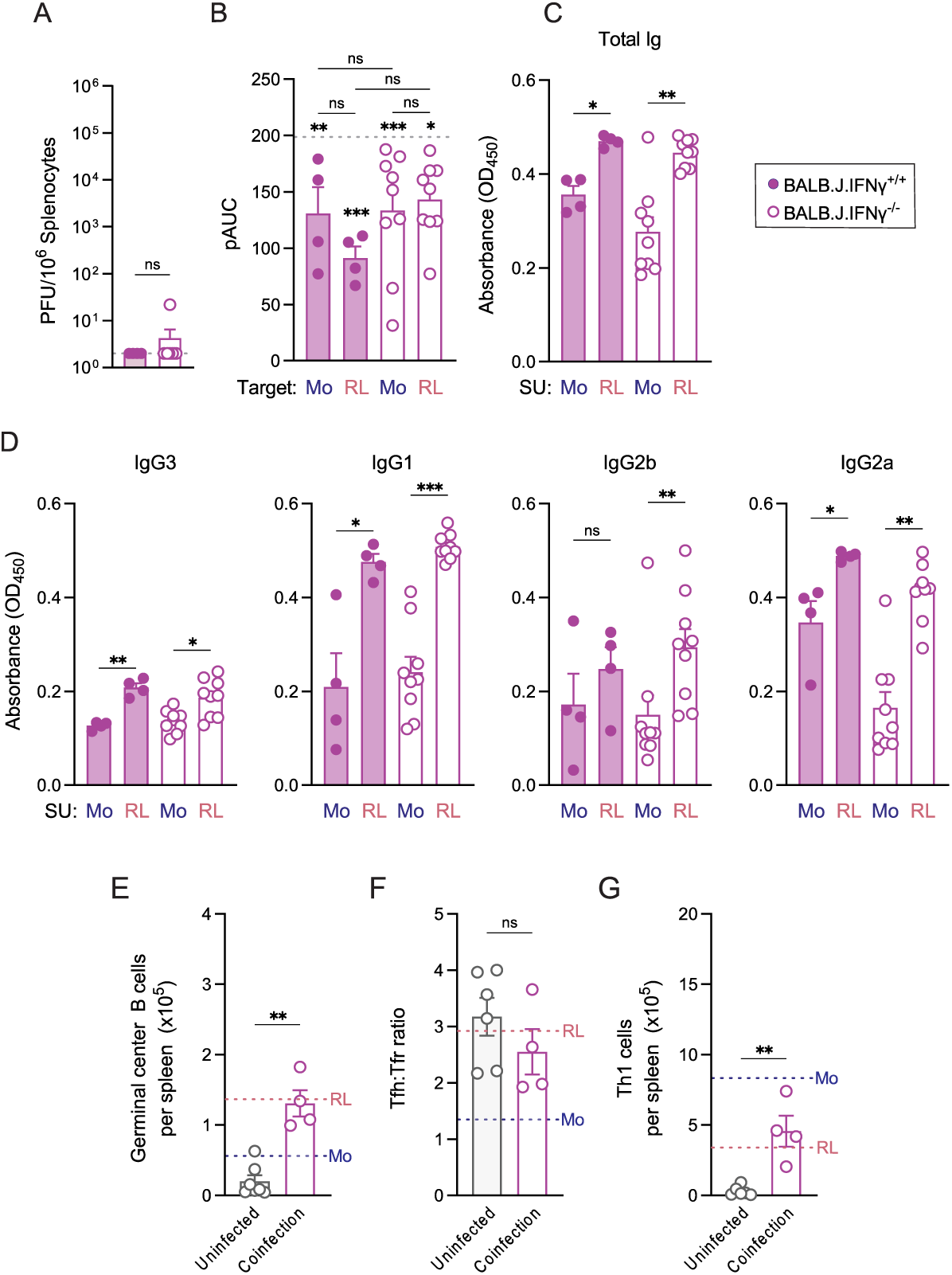
RL-MLV infection stimulates protective humoral immune responses. Mice of the indicated genotypes were infected with Mo-MLV and RL-MLV and analyzed eight weeks post infection. **(A)** Splenocytes were isolated and subjected to an infectious center assay. Dashed line indicates the limit of detection. **(B)** Sera was serially diluted and incubated with either Mo-MLV or RL-MLV before adding to SC-1 cells. Neutralization capacity was quantified by determining the partial area under the curve (pAUC). Grey dashed line represents the average pAUC of naïve sera. Statistics above columns represent comparisons against naïve sera. **(C)** Sera was collected and monitored for total Ig antibodies against MLV-SU protein. **(D)** Sera from infected mice were monitored for IgG3, IgG1, IgG2b, and IgG2a antibodies against Mo- or RL-SU by ELISA. **(E-G)** Splenocytes were isolated and analyzed by flow cytometry. Red dashed line represents mean of singular RL-MLV infected mice as reported in Fig. 2. Blue dashed line represents mean of singular Mo-MLV infected mice as reported in Fig. 2. **(E)** Number of germinal center (GC) B cells. **(F)** Ratio of Tfh:Tfr cells. **(G)** Number of T helper 1 (Th1) cells. *For each statistical comparison, a nonparametric Kruskal-Wallis test was applied, and corresponding significance values are indicated for each graph. ns, not significant; *, p<0.05; **, p<0.01; ***, p<0.001*.

### Gammaretroviral Env controls the capacity to induce IFNγ-independent antiviral immune responses

Next, we sought to establish the viral determinants for stimulation of IFNγ-independent antiviral immune responses. The MLV envelope glycoprotein mediates viral attachment and entry, is the target for neutralizing antibodies, and is the most divergent viral protein between Mo- and RL-MLV **(Fig. S3A)**. Thus, we generated chimeric viruses in which the *env* genes were exchanged between Mo- and RL-MLV **(Fig. 5A)**. IFNγ-deficient mice were infected with chimeric viruses and assessed for their antiviral antibody responses. We found that both IFNγ-sufficient and -deficient mice infected with chimeric Mo-MLV containing the RL-Env (Mo-MLV_RL-Env_) produced neutralizing antibodies and cleared the infection **(Fig. 5B-D)**. Additionally, IFNγ-deficient mice infected with the reciprocal chimera (RL-MLV_Mo-Env_) failed to generate neutralizing antibodies and retained infectious virus in their spleens **(Fig. 5B-D)**. These results indicate that RL-MLV Env is necessary and sufficient for the generation of neutralizing antibody responses in the absence of IFNγ. Next, we infected IFNγ-deficient BALB.J mice with chimeric viruses and assessed their B and T cell responses via flow cytometry. In IFNγ-deficient mice, Mo-MLV_RL-Env_ infection induced GC B cell responses and IgG2a class switching similar to RL-MLV infection, whereas RL-MLV_Mo-Env_ infection failed to generate a robust GC B cell response and GC B cells failed to class switch to IgG2a **(Fig. 5E-F)**. Furthermore, we found no difference in the number of Tfh cells between mice infected with chimeric viruses **(Fig. 5G)** but observed more Tfr cells in mice infected with RL-MLV_Mo-Env_ **(Fig. 5H)**, and thus the Tfh:Tfr ratio was skewed towards Tfr cells **(Fig. 5I)**. Finally, in both IFNγ-sufficient and -deficient mice, infection with RL-MLV_Mo-Env_ increased the number of Th1 cells **(Fig. 5J)**. Taken together, these results confirm that gammaretroviral Env is necessary and sufficient for the induction of protective IFNγ-independent antiviral immune responses. Next, to determine the domain of Env that controls IFNγ-independent antiviral antibody responses, we constructed SU chimeras by exchanging the SU between Mo- and RL-MLV **(Fig. 5K)**. IFNγ-deficient mice infected with chimeric Mo-MLV containing RL-SU produced neutralizing antibodies and cleared the infection **(Fig. 5L-N)**, while mice infected with RL-MLV containing Mo-SU exhibited an intermediate phenotype, since four of nine infected mice produced neutralizing antibodies **(Fig. 5M)** and cleared the infection **(Fig. 5N)**. The N-terminal portion of MLV-SU consists of three functional domains, the signal peptide (SP), receptor binding domain (RBD), and the proline-rich region (PRR) (Hogan and Johnson, 2023). All three domains contain amino acid differences between Mo- and RL-MLV **(Fig. S3B)**, making each a candidate for controlling antiviral antibody responses. The SP functions in trafficking and enhances packaging efficiency of viral particles (Liu et al., 2017), and the composition of the SP has been shown to impact glycosylation and antigenicity of the envelope glycoprotein (Yolitz et al., 2018). The RBD controls binding of the envelope glycoprotein to cellular receptors and is the target for neutralizing antibodies. The PRR is not well conserved between gammaretroviruses and functions as an antigenic decoy for non-neutralizing antibodies (Burkhart et al., 2003). As we have previously determined that antigenic differences between Mo- and RL-MLV do not determine neutralizing antibody responses, we sought to test whether RL-RBD stimulates protective immunity. We infected mice with Mo-MLV containing the RL-RBD and found that Mo-MLV_RL-RBD_ phenocopied Mo-MLV_RL-SU_ and RL-MLV **(Fig. 5L-N)**, demonstrating that RL-RBD is sufficient to stimulate IFNγ-independent neutralizing antibody responses.

**Figure 5.**
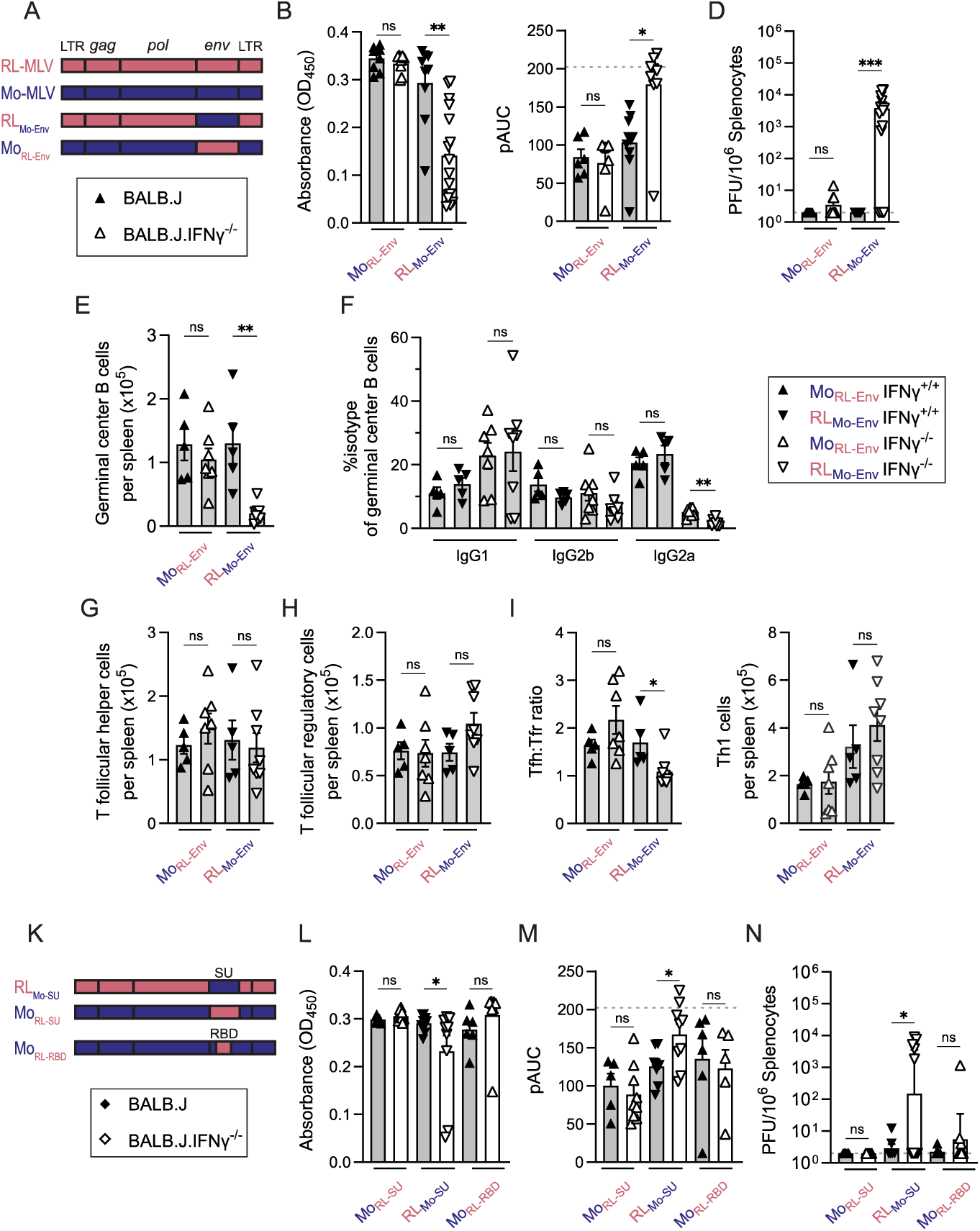
Gammaretroviral SU controls stimulation of IFNγ-independent neutralizing antiviral antibody responses. **(A)** Diagram of MLV chimeras. **(B-D)** Mice of the indicated genotypes were infected with MLV Env chimeras and analyzed eight weeks post infection. **(B)** Sera from infected mice were monitored for total Igs against respective MLV-SU protein by ELISA. **(C)** Sera was serially diluted and incubated with either RL-MLV or Mo-MLV before adding to SC-1 cells. Neutralization capacity was quantified by determining the pAUC. Grey dashed line represents the average pAUC of naïve sera. **(D)** Splenocytes were isolated and subjected to an infectious center assay. Grey dashed line indicates the limit of detection. **(E-J)** Mice of the indicated genotypes were infected with MLV Env chimeras and splenocytes were isolated and analyzed by flow cytometry four weeks post infection. **(E)** Number of germinal center (GC) B cells. **(F)** Frequency of GC B cells that were IgM^−^IgG1^+^, IgM^−^IgG2b^+^, and IgM^−^IgG2a^+^. **(G)** Number of T follicular helper (Tfh) cells. **(H)** Number of T follicular regulatory (Tfr) cells. **(I)** Ratio of Tfh to Tfr cells. **(J)** Number of T helper 1 (Th1) cells. **(K)** Diagram of SU and RBD chimeras with labeled viral genome. **(L-N)** Mice of the indicated genotypes were infected with MLV SU and RBD chimeras and analyzed eight weeks post infection. **(L)** Ig titers were determined as in (B). **(M)** Neutralization was determined as in (C). Grey dashed line represents the average pAUC of naïve sera. **(N)** Viral titers were determined as in (D). *For each statistical comparison, a nonparametric Kruskal-Wallis test was applied, and corresponding significance values are indicated for each graph. ns, not significant; *, p<0.05; **, p<0.01*.

### Sequence differences in gammaretroviral *env* do not control *in vitro* replication kinetics or *in vivo* viral spread

One possible explanation for Env controlling antiviral immune responses is that it alters infection efficiency resulting in differences in the magnitude of viral infection. To test this possibility, we first examined the *in vitro* replication kinetics of both Mo-MLV and RL-MLV and their respective *env* chimeras. Both Mo- and RL-MLV replicated at similar rates *in vitro* **(Fig. S4A)** and swapping the *env* genes did not affect replication **(Fig. S4A)**. To test whether *in vivo* viral spread differs between MLV strains, we infected both IFNγ-sufficient and -deficient BALB.J mice with Mo- and RL-MLV and measured viral titers in both bone marrow and splenic tissue over time. At one-week post-infection, RL-MLV infected mice had detectable PFUs in both the bone marrow **(Fig. S4B)** and spleen **(Fig. S4C)**, while we did not detect significant viral titers in Mo-MLV infected mice until two weeks post infection **(Fig. S4B-C)**. To determine whether these differences were controlled by Env, we repeated these experiments with *env* chimeric viruses and observed that mice infected with Mo-MLV_RL-Env_ phenocopied those infected with Mo-MLV, while mice infected with RL-MLV_Mo-Env_ phenocopied those infected with RL-MLV **(Fig. S4B-C)**. Thus, these results indicate that viral determinants outside of Env determine *in vivo* viral spreading kinetics, and that early viral spread does not determine the ability to stimulate neutralizing immune responses.

### MLV-SUs exhibit distinct binding affinities to various cell types, and RL-SU controls infection of Gr-1^+^ myeloid cells

All ecotropic strains of MLV share a common ubiquitously expressed receptor, mouse cationic amino acid transporter 1 (mCAT1); however, expression and glycosylation of mCAT1 varies among different cell types (Suzuki et al., 2001; Wang et al., 1996) and single residue changes in the SU and differences in receptor affinity have been shown to affect MLV tropism and pathogenesis (Masuda et al., 1996; Murphy et al., 2006). Therefore, a potential explanation for how SU controls the stimulation of neutralizing IFNγ-independent antibodies is that sequence differences in SU determine fusion and entry into key cell types that drive innate sensing of MLV, thus activating protective antibody responses. To test the binding affinity of Mo- and RL-SU to various hematopoietic and lymphoid stromal cells, we incubated recombinant SU proteins with a human IgG1-Fc tag with splenocytes and bone marrow (BM) cells isolated from naïve IFNγ-deficient BALB.J mice. Cells were then stained with a panel of antibodies for the identification of various hematopoietic and stromal cell types and SU binding was measured with a secondary antibody against human IgG1-Fc **(Fig. S5)**. To analyze which cells are infected by RL- and Mo-MLV *in vivo*, we infected IFNγ-deficient BALB.J mice with mNeonGreen reporter MLV (RL-MLV_mNG_ and Mo-MLV_mNG_), isolated splenocytes at two weeks post infection, and identified infected cells by flow cytometry. Previous reports have identified dendritic cells (DCs) and B cells as major targets of MLV infection (Pi et al., 2019; Podschwadt et al., 2022; Salas-Briceno et al., 2024); thus, we reasoned that differential SU binding and infection of DCs or B cells could explain the distinct outcomes of RL- and Mo-MLV infection. We observed that RL-SU bound to DCs more than Mo-SU **(Fig. 6A)**; however, binding affinity did not correlate with differences in infection of DCs **(Fig. 6B)**. Furthermore, we found that Mo-SU exhibited higher binding to B cells in comparison to RL-SU **(Fig. 6C),** but we observed no difference in frequency of infected B cells **(Fig. 6D)**. Importantly, we have previously demonstrated that the radiation-resistant compartment controls the IFNγ-independent antibody response (Kane et al., 2018). Since follicular dendritic cells (FDCs) are part of the radiation-resistant compartment and are critical to B cell responses, we investigated whether there were differences in binding or infection of FDCs by Mo- or RL-MLV. Mo-SU exhibited increased binding to FDCs compared to RL-SU **(Fig. 6E)**; however, there was no difference in frequency of infected FDCs **(Fig. 6F)**. Indeed, there was no overall correlation between SU binding affinity and infectivity. Furthermore, among all cell types examined, only Gr-1^+^ myeloid cells were infected at different levels by the two strains **(Figs. 6 and S7)**; and this difference was determined by SU, as shown by RL-SU binding and RL-MLV_mNG_ and Mo-MLV_RL-SU_,_mNG_ infecting Gr-1^+^ myeloid cells at higher levels than Mo-SU and Mo-MLV_mNG_ **(Fig. 6G-H)**. Taken together, these data demonstrate that MLV-SU variants differ in their binding affinity to various cell types and that RL-SU specifically determines the capacity to infect Gr-1^+^ myeloid cells.

**Figure 6.**
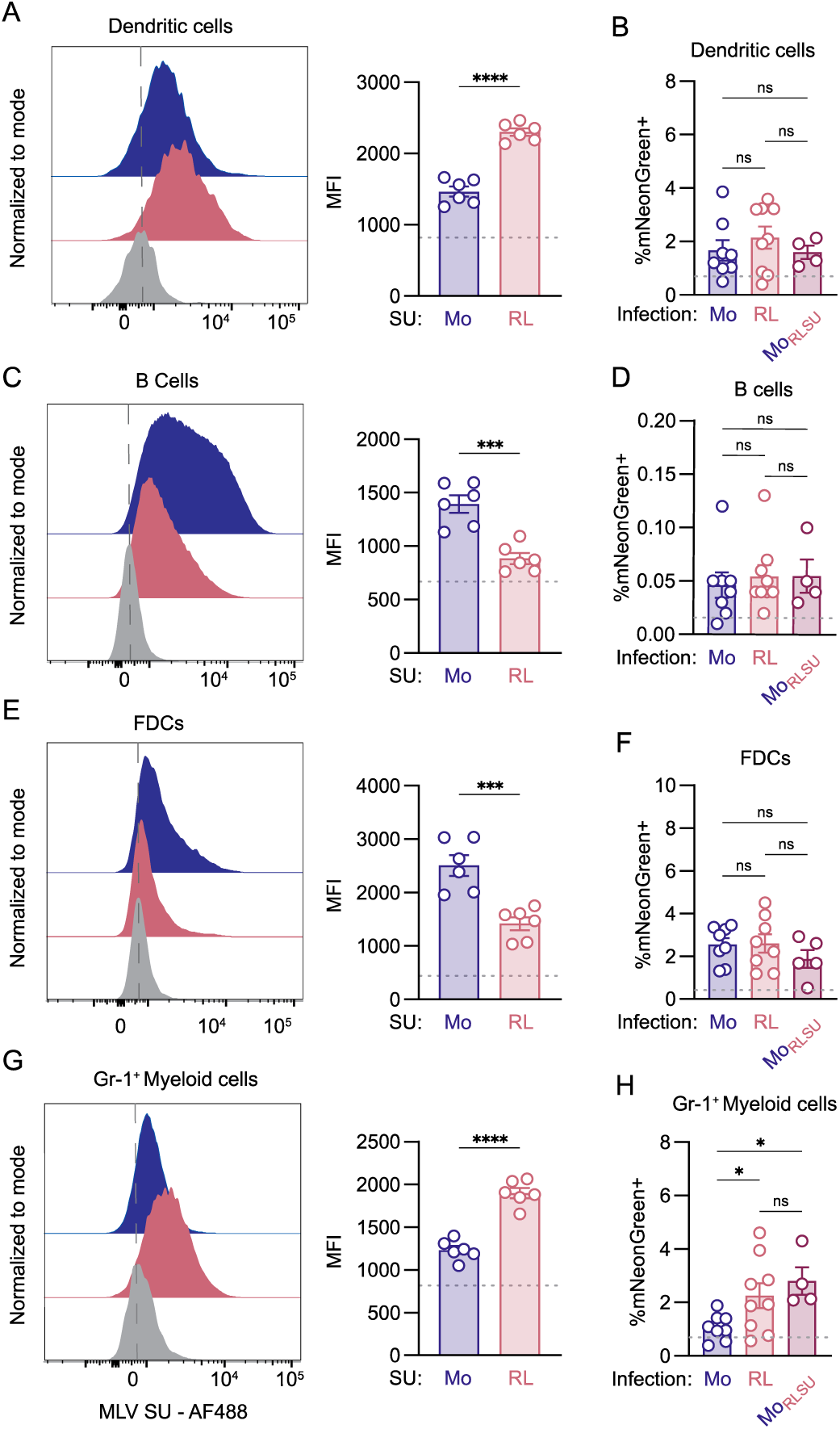
Mo- and RL-SU have distinct interactions with various cell types and SU determines infection of Gr-1^+^ myeloid cells. **(A,C,E,G)** Splenocytes were isolated from naïve IFNγ-deficient BALB.J mice and incubated with recombinant MLV-SU tagged with human IgG1-Fc, which was detected with anti-huIgG1 conjugated with Alexa Fluor 488 and analyzed by flow cytometry. **(B,D,F,H)** IFNγ-deficient BALB.J mice were infected with the indicated mNeonGreen reporter virus and analyzed by FACS. **(A)** Representative offset histogram plots for Gr-1^+^ myeloid cells and calculated MFI of the indicated MLV-SU. Grey dashed line indicates no SU. **(B)** Frequency of mNeonGreen positive Gr-1^+^ myeloid cells. **(C)** Representative offset histogram plots for bone marrow derived dendritic cells (BM-DC) and calculated MFI of the indicated MLV-SU. Grey dashed line indicates no SU. **(D)** Frequency of mNeonGreen positive BM-DC. **(E)** Representative offset histogram plots for B cells and calculated MFI of indicated MLV-SU. Grey dashed line indicates no SU. **(F)** Frequency of mNeonGreen positive B cells. **(G)** Representative offset histogram plots for FDC and calculated MFI of indicated MLV-SU. Grey dashed line indicates no SU. **(H)** Frequency of mNeonGreen positive FDC. *For two-group comparisons a Mann-Whitney test was applied, for multi-group comparisons a nonparametric Kruskal-Wallis test was applied, and corresponding significance values are indicated for each graph. ns, not significant; *, p<0.05; ***, p<0.001; ****, p<0.0001*.

## DISCUSSION

The viral determinants responsible for stimulation of protective immune responses are poorly defined. Here, we demonstrate that gammaretroviral Env, and specifically the RBD, determines the capacity to stimulate production of neutralizing antibodies in the absence of IFNγ signaling. We linked the extracellular subunit of the retroviral glycoprotein to differences in binding affinities to various cell types and infection of Gr-1^+^ myeloid cells. Thus, we hypothesize that the critical cell type controlling noncanonical antibody responses may be sensing RL-but not Mo-MLV and providing immunostimulatory signals upon RL-MLV infection. We observed that RL-SU bound more to Gr-1^+^ myeloid cells and BM-DCs, while Mo-SU exhibited higher binding to B cells and FDCs. Additionally, we found that viruses containing RL-SU infected a higher frequency of Gr-1^+^ myeloid cells than viruses with Mo-SU. One possibility is that productive infection of Gr-1^+^ myeloid cells is responsible for activating protective humoral immunity in the absence of IFNγ-signaling. Gr-1^+^ myeloid cells comprise a heterogeneous population of immature myeloid cells and are also known as myeloid-derived suppressor cells (MDSC) (Zanghi et al., 2024). MDSCs impair host immunity by preventing T-cell activation, secreting IL-10, and producing TGFβ and reactive oxygen species (Gabrilovich et al., 2012). Notably, in the context of the tumor microenvironment, stimulation of TLR7 signaling in the MDSC population removed immunosuppressive abilities, and the MDSC population acquired an antigen-presenting phenotype (Spinetti et al., 2016). Previous work by us and others has shown that TLR7-dependent sensing of retroviruses is required for activation of humoral immunity (Browne, 2011; Kane et al., 2011). Therefore, RL-MLV infection may stimulate protective immune responses upon infection of the MDSC population and subsequent induction of an immunostimulatory environment. Alternatively, detection of non-productive MLV infection in DCs may control IFNγ-independent antibody responses. Importantly, TLR signaling in plasmacytoid dendritic cells (pDCs) leads to the synthesis and secretion of type I and type III IFNs, and pDC-derived IFNs have been shown to be critical for control of coronavirus, LCMV, HSV, and influenza infections (Cervantes-Barragan et al., 2012; Cervantes-Barragan et al., 2007; Ciancanelli et al., 2015; Coccia et al., 2004; Swiecki et al., 2013). Furthermore, TLR signaling in immature conventional dendritic cells (cDCs) results in cDC maturation and migration to secondary lymphoid organs, where mature cDCs secrete IL-6 and IL-21 to promote Tfh maturation, thus supporting the GC B cell response (Durand et al., 2019; Krishnaswamy et al., 2017). We observed that RL-SU bound to DCs more efficiently than Mo-SU *in vitro*, but viruses containing RL-SU did not exhibit higher levels of infection in DCs *in vivo*; therefore, abortive infection of RL-MLV in DCs could stimulate TLR7 signaling and drive antiviral cytokine responses. Future experiments to elucidate the role of MDSC or DC populations may reveal the critical cell types required in the noncanonical pathway for neutralizing antibodies. The consideration of cell types stimulated by vaccination strategies could also inform the future development of therapeutics to prevent viral pathogenesis.

The importance of IgG2a antibodies in protection against viral infections in murine models is well established (Case et al., 2008; Case et al., 2005; Coutelier et al., 1987; Kane et al., 2018; Markine-Goriaynoff et al., 2000; McKendall and Woo, 1988; van den Broek et al., 1995; Zhang et al., 2023). In this work, we report that both the canonical IFNγ-dependent and noncanonical IFNγ-independent antiviral antibody response in BALB.J and 129S7 mice involve the generation of antiviral antibodies of multiple IgG isotypes (including IgG2a), and that closely related viruses can induce distinct antibody isotypes upon infection, suggesting that the previously observed bias towards IgG2a production may not be generally representative of antiviral immune responses. Additionally, earlier reports have demonstrated that immunization with viral proteins or peptides leads to a predominantly IgG1 antibody response (Balkovic et al., 1987; Purdy et al., 2003; Sallberg et al., 1996; Smucny et al., 1995), similar to responses to other protein antigens, suggesting that antigenic differences may not be responsible for modulation of the antibody response. However, other reports have indicated that immunization with inactivated viruses stimulates robust IgG1 and IgG2a responses (Balkovic et al., 1987; Deck et al., 1997; Markine-Goriaynoff et al., 2000; Peterson et al., 1992). These contrasting observations could be due to variance in methodology of inactivating virions, delivery of adjuvants, and strains of mice used. However, our results with both IFNγ-sufficient and -deficient mice immunized with Mo- and RL-MLV virions indicated that immunization with similar MLV antigens stimulated comparable IgG1, IgG2b, and IgG2a responses, suggesting that antigenic differences between MLV strains do not influence CSR. Additionally, while we found that B and CD4^+^ T cell responses were distinct upon infection with RL- or Mo-MLV, these responses did not differ upon immunization with viral proteins. Together, these results indicate that the nature of the immune response is controlled by events triggered by the infectious process rather than the antigenic properties of viral proteins. Interestingly, antibodies generated by infection with either Mo- or RL-MLV were cross-reactive but not cross-neutralizing; however, coinfection with Mo- and RL-MLV generated antibodies that neutralized both Mo- and RL-MLV virions. Further, coinfection induced germinal center responses similar to RL-MLV infection alone, demonstrating that RL-MLV infection stimulates the immune processes required to generate neutralizing antibodies in the absence of IFNγ. These distinct pathways provide a tractable system for identification of signals that drive protection against viral infection in future studies.

While the canonical pathway for antiviral antibody production is dependent on IFNγ signaling (Finkelman et al., 1988; Snapper and Paul, 1987; Snapper et al., 1988), the cytokines required for the noncanonical pathway remain unknown. What, then, could be the alternative cytokine driving non-canonical antibody responses? Importantly, the type I IFN response has also been shown to enhance the IgG2c antibody response *in vivo* (Swanson et al., 2010), indicating that type I IFNs may be important for controlling production of protective IFNγ-independent antibodies. Additionally, *in vitro* work has demonstrated that the heterodimeric cytokine IL-27 induces T-bet expression in B cells and regulates CSR to IgG2a (Yoshimoto et al., 2004). Further investigations into the *in vivo* role of candidate cytokines during RL-MLV infection will reveal the specific cytokine(s) and pathways required for noncanonical antibody responses.

## MATERIALS AND METHODS

### Mice

BALB.J mice (Denzin et al., 2017) and BALB.J.IFN-γ^−/−^ mice (Kane et al., 2018) were described previously and bred and maintained at the University of Pittsburgh. 129S1/SvImJ (The Jackson Laboratory, stock number 002448) (Stevens, 1973; Timmermans et al., 2017), 129-*Ifngr1^tm1Agt^*/J [also known as G129 mice (The Jackson Laboratory, stock number 002702)] (Kamijo et al., 1993), were bred and maintained at the University of Pittsburgh. Mice of both sexes were used in equal ratios for all experiments. All animal experiments were performed in the American Association for the Accreditation of Laboratory Animal Care-accredited, specific-pathogen-free facility at the Division of Laboratory Animal Resources, University of Pittsburgh School of Medicine. Animal protocols were reviewed and approved by the Institutional Animal Care and Use Committee at the University of Pittsburgh.

### Cell lines

SC-1 embryonic mouse fibroblasts (ATCC CRL-1404) and vir6 cells [SC-1 cells stably infected with the RL-MLV mixture (Hook et al., 2002)] were maintained in Dulbecco’s Modified Eagle Medium [(DMEM) Gibco] with 5% fetal calf serum [(FCS) Gibco] and gentamicin (Gibco). XC (ATCC CCL-165) were maintained in Modified Eagle Medium [MEM (Gibco)] with 10% FCS, sodium pyruvate (Gibco) and gentamicin. NIH3T3 cells (ATCC CRL-1658) were maintained in DMEM + 10% bovine calf serum (BCS) and gentamicin. HEK293T cells (ATCC CRL-3216) were maintained in DMEM with 10% FCS and gentamicin. 293F cells were maintained at 37°C with 8% CO_2_ in FreeStyle 293 Expression Medium (ThermoFisher) supplemented with penicillin/streptomycin (ThermoFisher). Mycoplasma testing (Invivogen) was regularly conducted on all cell lines, and cells were maintained in MycoZap prophylactic (Lonza) to prevent mycoplasma contamination. Prior to experimental use of cells, medium was changed to remove prophylactic.

### Plasmids and cloning

To determine the full sequence of ecotropic RL-MLV, viral RNA was isolated from virions purified from vir6 cells (Hook et al., 2002) via ultracentrifugation on a 30% sucrose gradient using a NucleoSpin RNA Virus kit (Macherey Nagel). First-strand cDNA was synthesized using SuperScript III (Invitrogen), followed by PCR amplification of overlapping pieces with oligos based on the Rauscher MLV sequence (GenBank: U94692.1) [**Table S1** Piece 1 (5’ R to Pol): R MLV F ‘R MLV 5514; Piece 4 (Pol to 3’ R): R MLV 5038 F + R MLV 8280R], and TOPO cloning (ZeroBlunt TOPO, Invitrogen). Four clones of each piece were sequenced. To generate the infectious molecular clone, pSRL, RL-MLV sequence was synthesized as GeneBlocks and cloned into pNCS (Yueh and Goff, 2003) using *Sfi*I and *Nde*I replacing the Mo-MLV sequence. Block 1: *Sfi*I-pNCS vector-*Not*I-5’ LTR-*gag*-*Hind*III; Block 2: *Hind*III-*gag-pol*,-*Sal*I; Block 3: *Sal*I-*pol-env*-*Age*I; Block 4: *Age*I-*env*-3’LTR-*Xho*I-*Nde*I. The *Nde*I and *Xho*I sites following the 3’-LTR were then replaced with an *EcoR*I site.

The RL-MLV_Mo-Env_ chimera was generated by swapping the Mo Env sequence from pNCS (Yueh and Goff, 2003) for the RL Env in pSRL. The sequence between *Nde*I (located in *pol*) and the *EcoRI* site (located in the LTR) was synthesized as a gBlock fragment (IDT) and subcloned using restriction enzymes *Nde*I and *EcoR*I (NEB). The RL-MLV_Mo-SU_ chimera was generated by swapping the Mo-SU sequence for the RL-SU in pSRL. The sequence between *Nde*I (located in *pol*) and the *EcoR*I site (located in the LTR) was synthesized as a gBlock fragment (IDT) and subcloned using restriction enzymes *Nde*I and *EcoR*I (NEB). The Mo-MLV_RL-Env_ chimera was generated by swapping the RL Env sequence for the Mo Env in pNCS. The sequence between *Nde*I (located in *pol*) and the *Nhe*I site (located in the LTR) was synthesized as a gBlock fragment (IDT) and subcloned using restriction enzymes *Nde*I and *Nhe*I (NEB). The Mo-MLV_RL-SU_ chimera was generated by swapping the RL-SU sequence for the Mo-SU in pNCS. The sequence between *Nde*I (located in *pol*) and the *Nhe*I site (located in the LTR) was synthesized as a gBlock fragment (IDT) and subcloned using restriction enzymes *Nde*I and *Nhe*I (NEB). The Mo-MLV_RL-RBD_ chimera was generated by swapping the RL-RBD sequence for the Mo-RBD in pNCS. The sequence between *Nde*I (located in *pol*) and the *Nhe*I site (located in the LTR) was synthesized as a gBlock fragment (IDT) and subcloned using restriction enzymes *Nde*I and *Nhe*I (NEB). The RL-MLV-mNeonGreen reporter construct was generated by inserting a P2A sequence followed by the mNeonGreen sequence between the 3’ end of *env* and the long terminal repeat (LTR) in the infectious plasmid clone of RL-MLV, pSRL. The sequence between the *Nde*I site (located in *pol*) and the *EcoR*I site (located in the LTR), including the P2A and mNeonGreen sequences, was synthesized as a gBlock fragment (Azenta) and subcloned into pSRL using restriction enzymes *Nde*I and *EcoR*I (NEB). The Mo-MLV-mNeonGreen reporter construct was generated by inserting a P2A sequence followed by the mNeonGreen sequence between the 3’ end of *env* and the long terminal repeat (LTR) in the infectious plasmid clone of Mo-MLV, pNCS. The sequence between the *Nde*I site (located in *pol*) and the *Nhe*I site (located in the LTR), including the P2A and mNeonGreen sequences, was synthesized as a gBlock fragment (Azenta) and subcloned into pNCS using restriction enzymes *Nde*I and *Nhe*I (NEB).

The Mo-MLV_RL-SU,mNG_ reporter construct was generated using Gibson Assembly® Master Mix according to manufacturer’s instructions (NEB). In brief, DNA products were assembled with PCR products from Mo-MLV_RL-SU_ and Mo-MLV_mNG_ and with a *Nde*I (NEB) digested backbone from Mo-MLV_mNG_. The PCR product from Mo-MLV_RL-SU_ was amplified from the *NdeI* site (located in *pol*) and the 3’ end of *env*. The PCR product from Mo-MLV_mNG_ was amplified from the 3’ end of *env* to the *Nde*I site (located at the beginning of the LTR).

The full nucleotide sequence encoding amino acids 1-475 of the RL-SU domain was amplified from pSRL using PCR. Primers added a 5’ *Sal*I site and a GCCACC Kozak sequence before the open reading frame and a 3’ *Kas*I site using Phusion High-Fidelity DNA Polymerase (NEB) and primers KRM 597 and KRM 598. Amplicons were digested with *Sal*I and *Kas*I and cloned into a modified pVRC8400 vector encoding for a Gly-Ala linker, a 3C protease recognition site and a 6xHis tag (Robinson-McCarthy et al., 2018) which was linearized using the same enzymes. RL-SU was further modified by mutating its CXXC motif to SXXS using QuickChange site-directed mutagenesis (Agilent) and primers KRM 611 and KRM 612. The homologous region of Mo-MLV (amino acids 1-465) was cloned and modified using the same approach using pNCS as the template, primers KRM 595 and KRM 596 for amplification and KRM 605 and KRM 606 for mutagenesis. This expression construct did not yield measurable quantities of protein. Mo-SU was cloned into a modified pVRC8400 vector encoding for a 3’ Gly-Ala linker, a human IgG1 Fc domain, a Gly-Gly linker and a 6xHis tag using *Sal*I and *Kas*I sites (NEB). The modified RL-MLV SU was similarly cloned into this expression vector.

### Virus production

Stocks of replication competent ecotropic MLV strains (RL-MLV, Mo-MLV, RL-MLV_Mo-Env_, Mo-MLV_RL-Env_, RL-MLV_Mo-SU_, Mo-MLV_RL-SU_, Mo-MLV_RL-RBD_) used for infecting mice were generated by transfecting NIH3T3 cells with 10 μg of proviral plasmid DNA using Lipofectamine® LTX Reagent (Thermofisher) according to manufacturer’s protocol. Replication competent ecotropic MLV was titered by infectious center assay.

Ecotropic mNeonGreen reporter viruses used for infecting mice (Mo-MLV_mNG_, RL-MLV_mNG_, and Mo-MLV_RL-SU,mNG_) were generated by Neon NxT Electroporation of NIH3T3 cells and concentrated with Lenti-X^TM^ Concentrator (TaKaRa) according to manufacturer’s protocol and resuspended in PBS. Reporter viruses were titered by infection of SC-1 cells and enumerated by FACS analysis. Ecotropic reporter viruses used for neutralization assays were generated by transfecting HEK293T cells using PEI (PolySciences) for 48-72 hours. Harvested supernatant was filtered, and mNeonGreen reporter virus stocks were titered by serial dilution and incubation with SC-1 cells for 48 hours.

### *In vitro* spreading assay

For *in vitro* spreading, viral supernatants were quantified using the SYBR-Green based PCR reverse transcription (RT) Assay (Vermeire et al., 2012). In brief, 2.5×10^3^ PFU of each virus was used to infect 2.5×10^4^ SC-1 cells. 16h post infection, cells were washed and supernatants were collected over 14 days. 5μl of cell culture supernatant was incubated with 5μl 2x lysis buffer (0.25% Triton X-100, 50mM KCl, 100mM TrisHCl pH7.4, 40% glycerol) for 10 min at RT. The lysate was diluted 10x in 1x Core buffer: 5 mM (NH4)2SO4, 20 mM KCl and 20 mM Tris–HCl pH 8.3. 10μl of the sample were then mixed with 10μl of 2x reaction buffer: 5mM (NH4)2SO4, 20mM KCl and 20mM Tris–Cl (pH8.3), 10mM MgCl2, 0.2mg/ml BSA, 1x dilution of SYBR Green I (Life technologies S-7563), 400μM dNTPs, 1μM MS2-F oligo, 1μM MS2-R oligo, 0.0002U/ml MS2 RNA (USBiological Life Sciences). RT reaction conditions were 42°C for 20 min, 95°C for 2 min, followed by 40 cycles of 95°C for 5 sec, 60°C for 5 sec, 72°C for 15 sec and 80°C for 10 seconds measured using the QuantStudio 3 Real-Time PCR System (Thermofisher). The RT assay was also used to quantify RT levels in viral stocks.

### Infectious center assay

MLV viral titers were determined by infectious center assay (Rowe et al., 1970) as previously described (Zhang et al., 2023). In brief, irradiated splenocytes were incubated with SC-1 cells for 5 days. SC-1 cells were killed by UV-irradiation and overlaid with XC cells for 2 days. Cells were then stained with a mixture of methylene blue (Thermofisher) and carbol fuchsin (Sigma-Aldrich) in methanol, and localized syncytia were counted.

### Infection and immunization

For MLV infection, experimental mice were injected i.p. with 2 × 10^4^ ecotropic MLV PFUs (as determined by infectious center assay) at 5-8 weeks of age. For MLV immunization, experimental mice were injected i.p. with 50µg of Triton-X-100-treated MLV virion proteins mixed with Complete Freund’s Adjuvant (Invivogen) at 5-8 weeks of age. For mNeonGreen labeled MLV infection, experimental mice were injected i.p. with 2 × 10^5^ mNeonGreen labeled MLV at 5-8 weeks of age.

### Recombinant SU

Recombinant MLV SU and SU-Fc fusion proteins were expressed and purified as previously described (McCarthy et al., 2020; Simmons et al., 2023). Briefly, 293F cells were transiently transfected with SU or SU-Fc plasmids using PEI. Transfection complexes were prepared in Opti-MEM and added to cells. Supernatants were harvested 4–5 days post-transfection and clarified by low-speed centrifugation (3500xg). Proteins were purified by adsorption to cobalt-nitrilotriacetic acid (Co-NTA) agarose resin (Takara Bio USA), washed in 150mM sodium chloride, 10mM tris(hydroxymethyl)aminomethane-hydrochloride (pH 7.5) (buffer A). Protein was eluted in buffer A plus 350 mM imidazole (pH 8), concentrated in an ultra centrifugal filter with a 10 kDa molecular weight cutoff (Amicon) and further purified by gel filtration chromatography in buffer A on a Superdex 200 column (Cytiva). ELISA using mouse serum (detailed below) was used to verify that the addition of a human IgG1-Fc region did not impact antibody binding (Fig. S8).

### ELISA

To detect MLV antibodies in mouse sera, an enzyme-linked immunosorbent assay (ELISA) was performed as previously described (Zhang et al., 2023). Virions isolated from MLV infected SC-1 cells were treated with 0.1% Triton X-100 or purified MLV SU was bound to plastic in borate-buffered saline overnight at 250 ng per well, followed by incubation with mouse serum samples at 4°C for one hour. All sera were used at 2 × 10^−2^ dilutions. Mouse Ig-specific, secondary antibodies coupled to horseradish peroxidase (HRP; Jackson ImmunoResearch) were used to detect antivirus antibodies. Ovalbumin (2%) was used as a blocking reagent. Backgrounds obtained from incubation with secondary antibodies alone were subtracted from the values obtained from sera of infected mice.

### Neutralization

Sera from MLV-infected and control mice were tested for their ability to neutralize virus. mNeonGreen reporter RL- or Mo-MLV virions were incubated with serially diluted heat-inactivated sera from infected mice for one hour at room temperature. Virus/antibody mixtures were added to permissive SC-1 cells. 48 hours later, cells were trypsinized, fixed, and infected cells (%mNeonGreen positive) were determined by FACS analysis using an Attune NxT flow cytometer coupled to an autosampler. Neutralization capacity of serum was calculated using the partial area under the curve (Yu et al., 2012) and normalized to sera from naïve mice.

### Flow Cytometry

Splenocytes were isolated and made into single cell suspensions by mechanical disruption. Red blood cells were lysed with sterile distilled water (Gibco). White blood cells were washed and resuspended in PBS, and live cells were enumerated with trypan blue and Countess 3 Automated Cell Counter (Thermofisher). Cells were pelleted and washed once in PBS before staining. For all stains, two million cells per sample were stained with Zombie Aqua or Zombie NIR Fixable Viability Kit (BioLegend) according to manufacturer’s protocol and washed with FACS buffer [PBS (Corning) with 1% bovine serum albumin (Invitrogen), and 0.01% sodium azide)]. Cells were stained with fluorescently conjugated antibodies as labeled in figures for 30 min on ice, then washed with FACS buffer, and fixed with 2% PFA in PBS for 5 minutes on ice. For intracellular stains, cells were fixed/permeabilized with FoxP3 Transcription Factor staining Kit (Invitrogen) according to manufacturer’s protocol. After fixation, cells were stained with intracellular stain antibodies as labeled in figures overnight at 4°C. Cells were then washed three times with permeabilization buffer, resuspended in FACS buffer, and data was collected using a Cytek Aurora flow cytometer and analyzed with FlowJo software.

### Statistical analysis

We used nonparametric Mann-Whitney tests to analyze two-group comparisons. Multi-group comparisons were analyzed by nonparametric Kruskal-Wallis test. All statistical analyses were performed with GraphPad Prism 11, with significance defined as P < 0.05.

## Data Availability

Data are available in the article itself and its supplementary materials.

## Acknowledgements

We thank members of the Kane and Golovkina laboratories for helpful discussion and Steve Joachim and Rebecca Elsner for technical assistance with flow cytometry and helpful discussion. This work was supported by grants from the NIH: R03AI180680 (to M.K.), R35GM154844 (to K.R.M), and T32 AI049820 (R.Z.Z.). This work was also supported by the Children’s Hospital of Pittsburgh of the UPMC Health System and the Diane and Cliff Rowe Research Fund. The content is solely the responsibility of the authors.

