## Supplementary Information for "Determinants in gammaretroviral Env dictate the production of neutralizing antibodies"

**This PDF file includes:**

Figures S1 to S7
Table S1 to S2

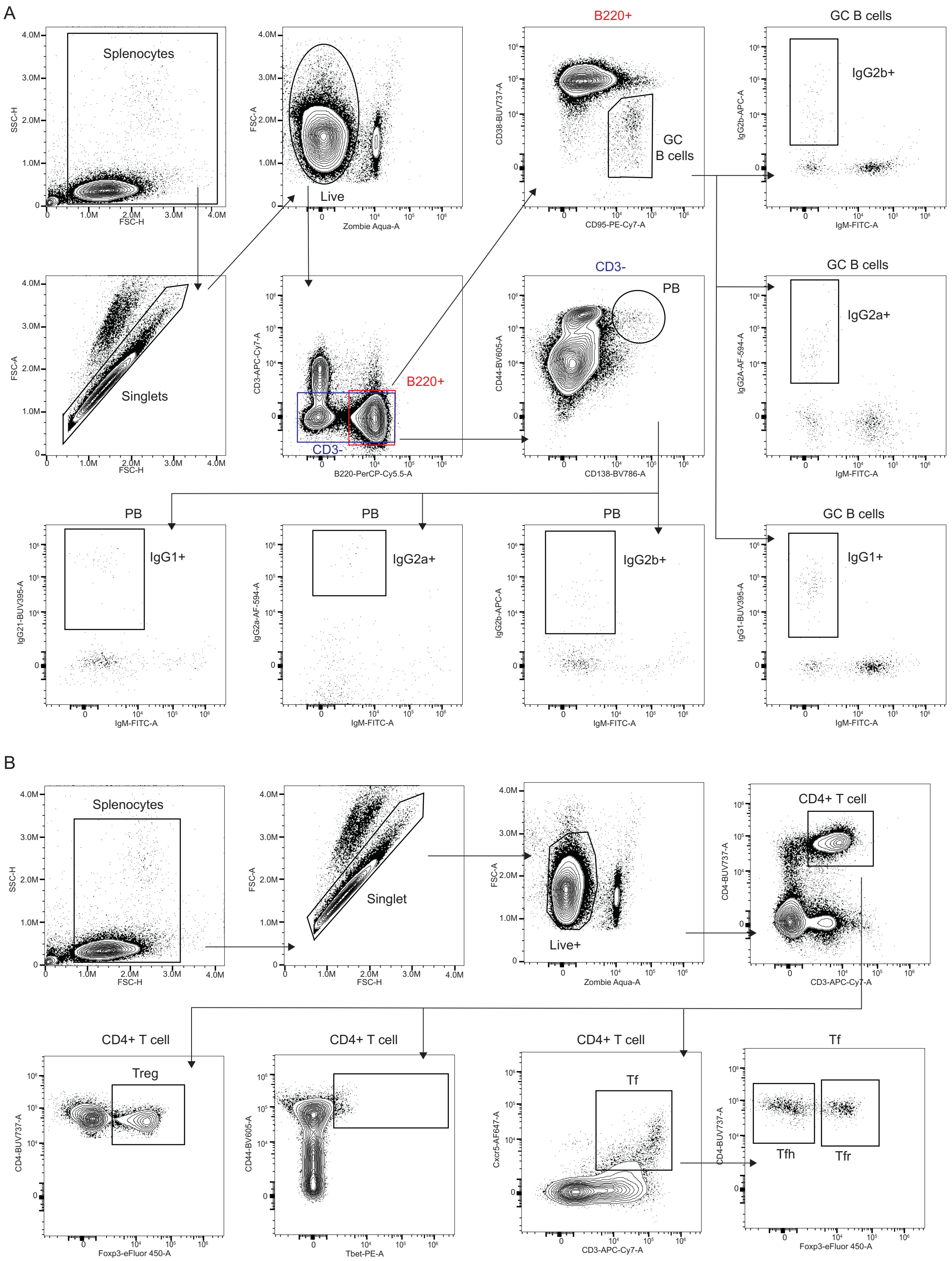


**Figure. S1. Gating strategies for lymphocyte subsets.** (A) Representative FACS plots of splenic GC B cells. (B) Representative FACS plots of various CD4+ T cell subsets in splenocytes.

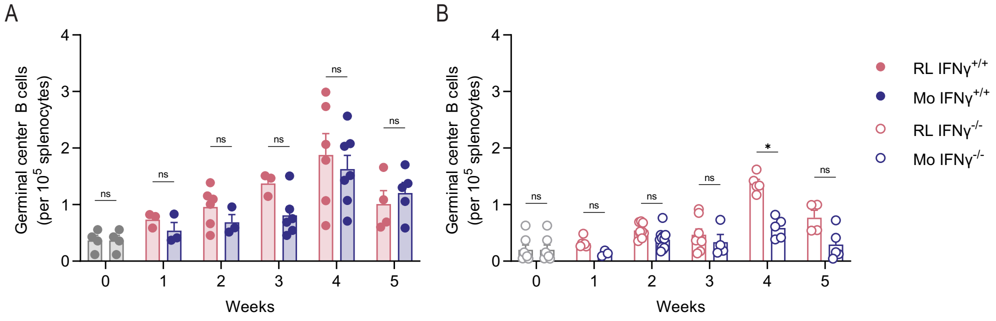


**Figure S2. IFNγ-deficient mice infected with Mo-MLV do not generate germinal center responses**. **(A)** BALB.J mice were infected with RL-MLV or Mo-MLV and monitored for frequency of GC B cells by FACS analysis at indicated timepoints. Gray dots represent naïve mice. **(B)** IFNγ-deficient BALB.J mice were infected with either RL-MLV or Mo-MLV and monitored for frequency of GC B cells by FACS at indicated timepoints. Open gray dots represent naïve mice. *For each statistical comparison, a nonparametric Mann-Whitney test was applied, and corresponding significance values are indicated for each graph.* ns, not significant; *, $p<0.05$.

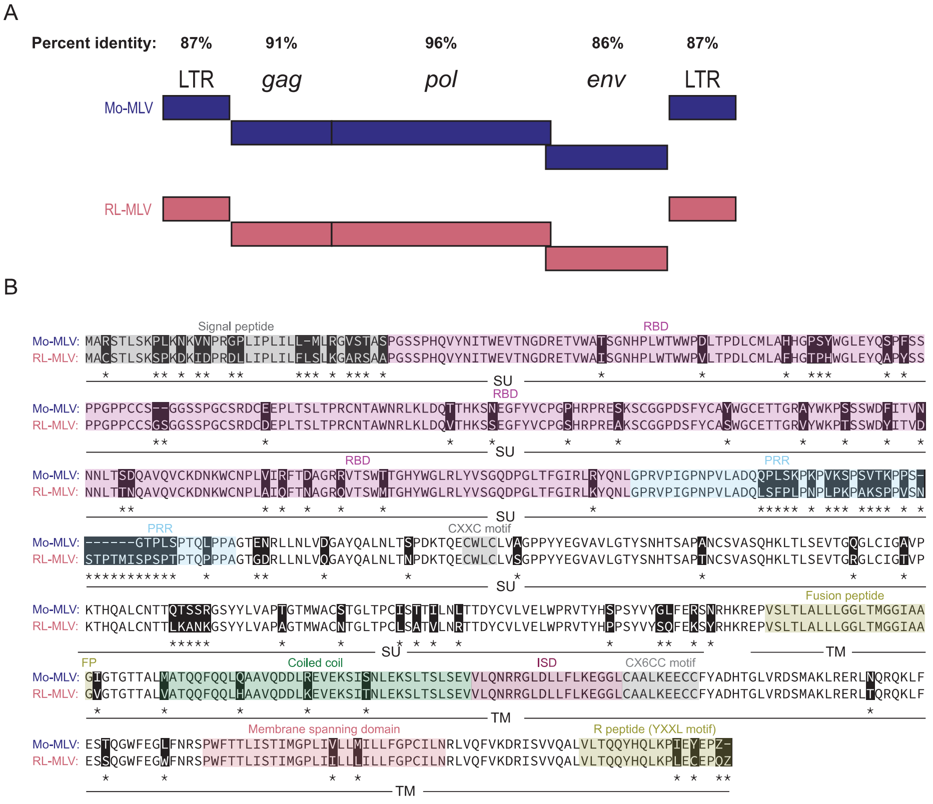


**Figure S3. Genome comparisons of Mo- and RL-MLV. (A)** Diagram of Mo-MLV genome compared to RL-MLV genome. Percent identity for LTR region is percent nucleotide similarity. Percent identity for *gag, pol,* and *env* is percent amino acid similarity. **(B)** Diagram of amino acid sequences for both Mo- and RL-MLV envelope glycoprotein with differences highlighted in black with an asterisk. Labeled are the surface unit (SU) and transmembrane unit (TM) as well as the domains.

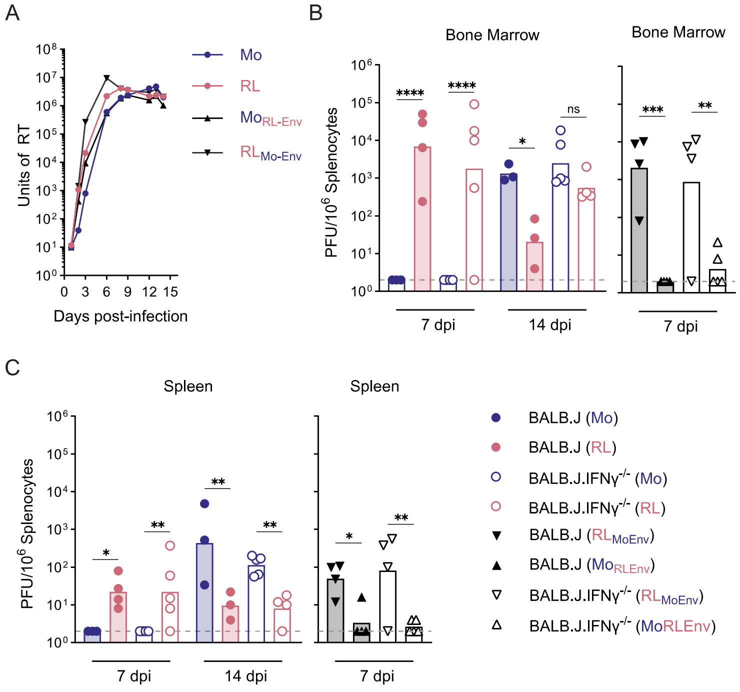


**Figure S4. Sequence differences in gammaretroviral Env do not control *in vitro* replication kinetics or *in vivo* viral spread. (A)** Spreading replication of RL-MLV, Mo-MLV, Mo-MLV$_{RL\text{-}Env}$, and RL-MLV$_{Mo\text{-}Env}$ (starting MOI = 0.01) in SC-1 cells. Infection was measured via SYBR-Green RT assay. **(B-C)** Mice of the indicated genotypes were infected with RL-MLV, Mo-MLV, Mo-MLV$_{RL\text{-}Env}$, or RL-MLV$_{Mo\text{-}Env}$. **(B)** Bone marrow cells were isolated and subjected to an infectious center assay at the indicated timepoints. Grey dashed line indicates limit of detection. **(C)** Splenocytes were isolated and subjected to an infectious center assay at indicated time points. Grey dashed line indicates limit of detection. *For each statistical comparison, a nonparametric Kruskal-Wallis test was applied, and corresponding significance values are indicated for each graph. ns, not significant; *, p<0.05; **, p<0.01; ****, p<0.0001*

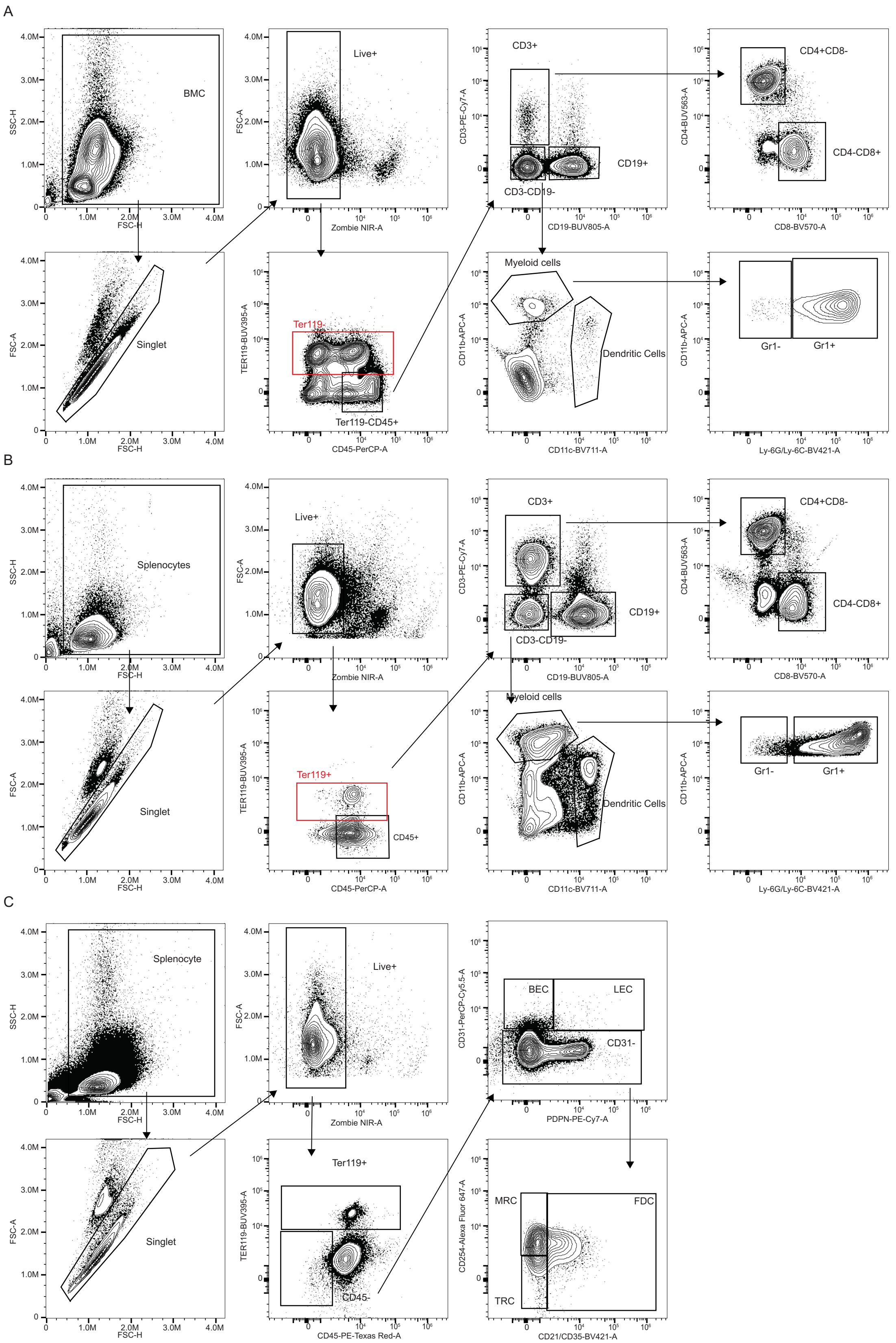


**Figure S5. Gating strategies for hematopoietic and non-hematopoietic cells. (A)** Representative FACS plots of hematopoietic cells from the bone marrow. **(B)** Representative FACS plots of hematopoietic cells from the spleen. **(C)** Representative FACS plots of stromal cells from the spleen.

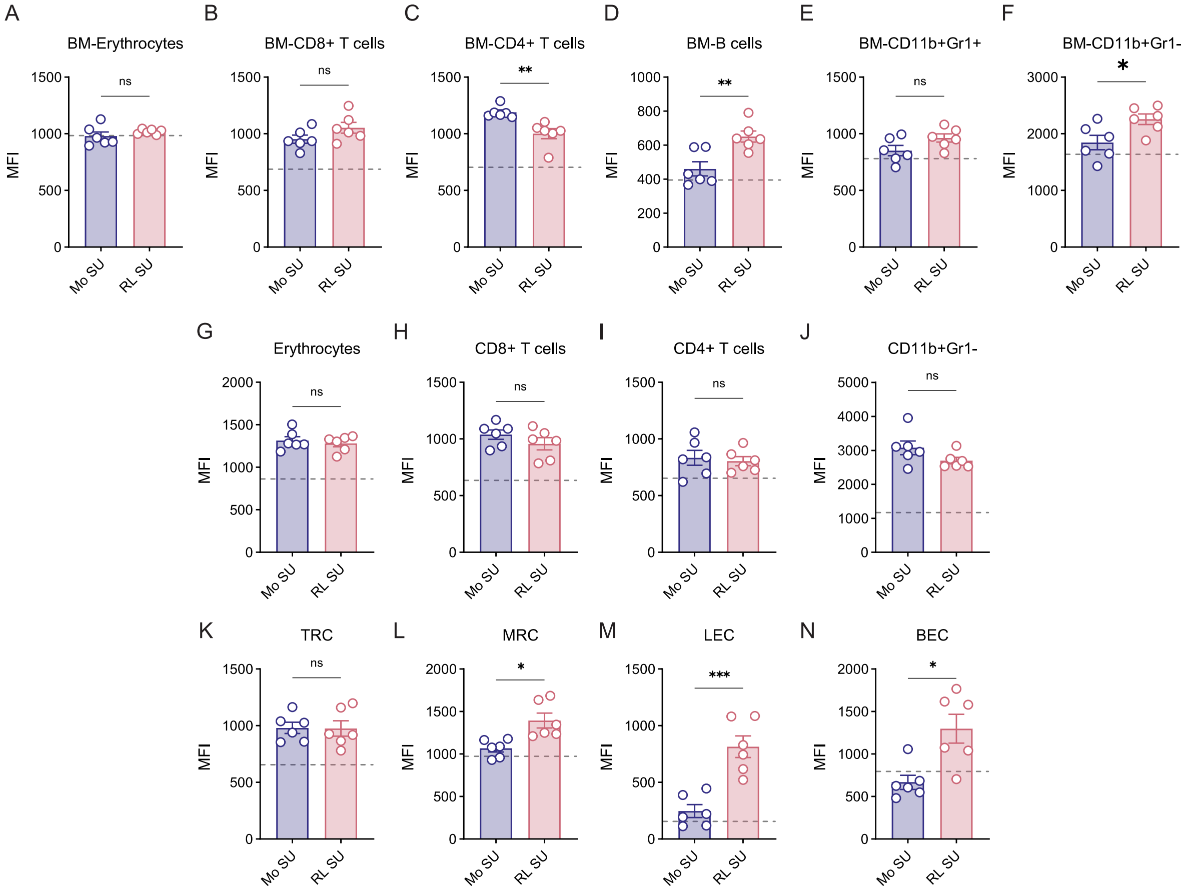


**Figure S6. Binding of Mo- and RL-SU to various cell types.** Splenocytes were isolated from naïve IFNγ-deficient BALB.J mice and incubated with recombinant MLV-SU tagged with human IgG1-Fc which was detected with anti-huIgG1 conjugated with Alexa Fluor 488 and analyzed by flow cytometry. Grey dashed line represents no SU control. **(A-F)** Cells isolated from bone marrow **(A)** Calculated MFI of MLV-SU in erythrocytes. **(B)** Calculated MFI of MLV-SU in $CD8^{+}$ T cells. **(C)** Calculated MFI of MLV-SU in $CD4^{+}$ T cells. **(D)** Calculated MFI of MLV-SU in B cells. **(E)** Calculated MFI of MLV-SU in $Gr\text{-}1^{+}$ myeloid cells. **(F)** Calculated MFI of MLV-SU in $Gr\text{-}1^{-}$ myeloid cells. **(G-N)** Cells isolated from spleen. **(G)** Calculated MFI of MLV-SU in erythrocytes. **(H)** Calculated MFI of MLV-SU in $CD8^{+}$ T cells. **(I)** Calculated MFI of MLV-SU in $CD4^{+}$ T cells. **(J)** Calculated MFI of MLV-SU in $Gr\text{-}1^{-}$ myeloid cells. **(K)** Calculated MFI of MLV-SU in T reticular cells. **(L)** Calculated MFI of MLV-SU in marginal reticular cells. **(M)** Calculated MFI of MLV-SU in lymphatic endothelial cells. **(N)** Calculated MFI of MLV-SU in blood endothelial cells. *For each statistical comparison, a nonparametric Mann-Whitney test was applied, and corresponding significance values are indicated for each graph.* ns, not significant; *, $p<0.05$; ***, $p<0.001$; ****, $p<0.0001$

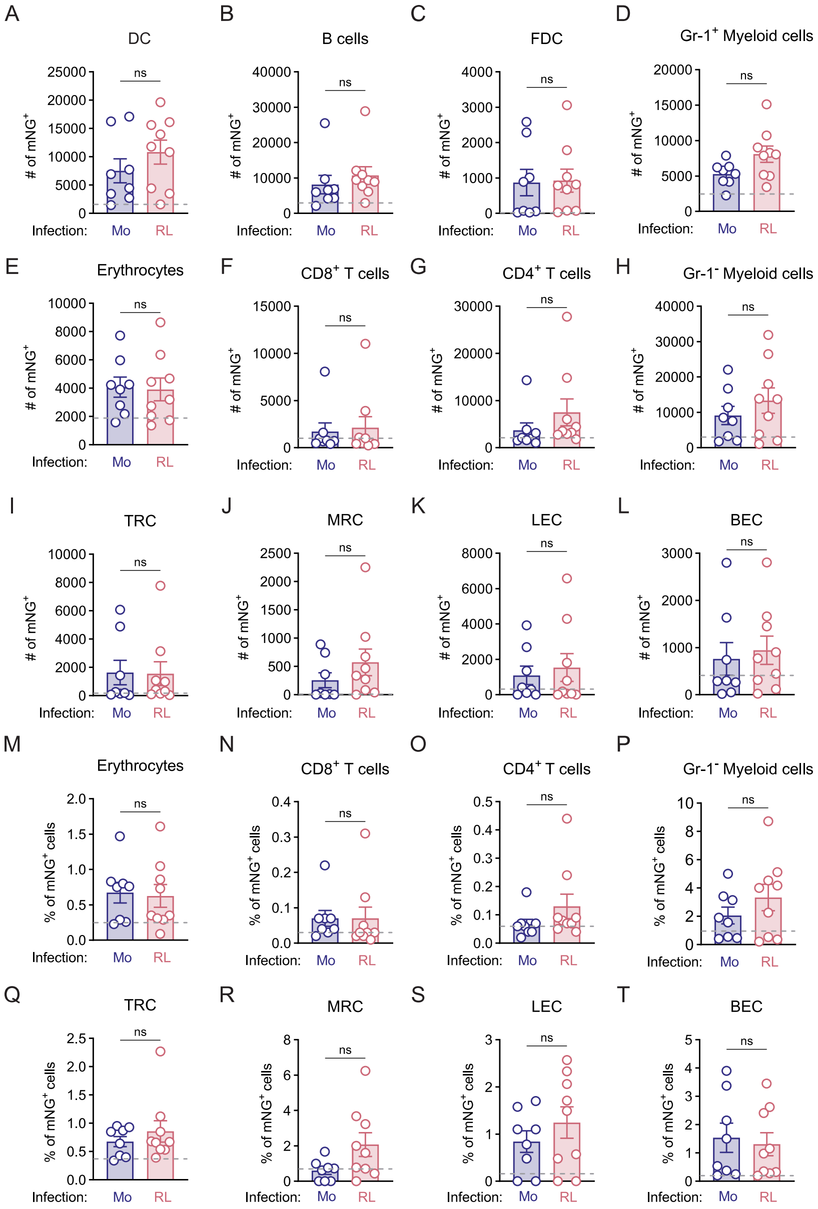


**Figure S7. Infection of Mo- and RL-MLV in various cell types.** Splenocytes were isolated from IFNγ-deficient BALB.J mice infected with Mo- and RL-MLV$_{mNG}$ viruses two weeks post infection and analyzed by flow cytometry. Grey dashed line represents splenocytes isolated from naïve mice. Number of NG$^+$ cells of total **(A)** dendritic cells **(B)** B cells **(C)** follicular dendritic cells **(D)** Gr-1$^+$ myeloid cells **(E)** erythrocytes **(F)** CD8$^+$ T cells **(G)** CD4$^+$ T cells **(H)** Gr-1$^-$ myeloid cells **(I)** T reticular cells and **(J)** marginal reticular cells **(K)** lymphatic endothelial cells **(L)** blood endothelial cells. Frequency of NG$^+$ cells of total **(M)** erythrocytes **(N)** CD8$^+$ T cells **(O)** CD4$^+$ T cells **(P)** Gr-1$^-$ myeloid cells **(Q)** T reticular cells and **(R)** marginal reticular cells **(S)** lymphatic endothelial cells and **(T)** blood endothelial cells. *For each statistical comparison, a nonparametric Mann-Whitney test was applied, and corresponding significance values are indicated for each graph.* ns, not significant.

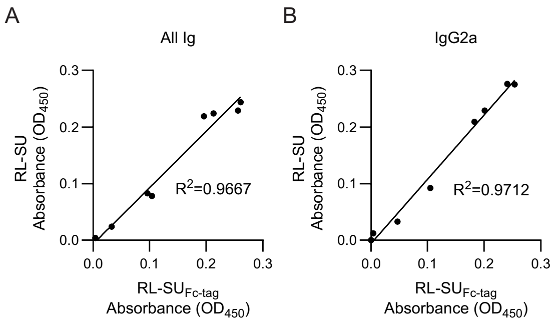


**Figure S8. Human IgG1-Fc Tag does not affect antibody binding to MLV-SU**. Representative sera samples were monitored for **(A)** total Igs and **(B)** IgG2a against recombinant RL-SU with and without human IgG1-Fc. *For statistical analysis, a simple linear regression was performed to determine the R-squared value.*

**S1 Table. Oligonucleotides for cloning and sequencing**

| | |
|---|---|
| RL-Pol-NdeI-F | CTCACCCCATATGAAATCTTATATGGGGCAC |
| RL-MLV 4-EcoRI R | TAACAGAGGAATTCAATGAAAGACCCCCGAGAC |
| Mo-Pol-NdeI-F | GAACTCTTTGTCGACGAGAAGCAGGG |
| Mo-NheI-LTR-R | GCGGTATTTCACACCGCATATGAAAGACCCCCGCTGACGG |
| Mo-Env-2A-F | AGCCTATAGAGTACGAGCCAGGTTCCGGAGCCACGAACTT |
| Mo-Env-2A-R | AAGTTCGTGGCTCCGGAACCTGGCTCGTACTCTATAGGCT |
| MS2-F | TCCTGCTCAACTTCCTGTCGAG |
| MS2-R | CACAGGTCAAACCTCCTAGGAATG |
| R MLV 2 F | GCCAGTCCTCCGACAGACTG |
| R MLV 658 F | GACCACTGGAAAGATGTCGAAC |
| R MLV 982 F | CTCTCGACCCCGCCTCAATCC |
| R MLV 1389 R | CTCGATCAAAGCTGTCAATTTACC |
| R MLV 1761 R | GGAGTGTATCTGCGATAGGCCTC |
| R MLV 2348 F | TCCTGGACCCCTAAGTGACAAG |
| R MLV 2635 R | AGGTCTCATGTAGCCGATACTCATC |
| R MLV 3702 F | GTTGACGAGAAGCAGGGCTAC |
| R MLV 3898 R | AATGACTAGTGGCTGTCCCATGG |
| R MLV 5038 F | GACACCTTCTCTGGATGGATAGAAGC |
| R MLV 5514 R | GACCAGGTAGAGTGCCTGTAAAT |
| R MLV 5589 F | CCGAGTCGGTGACACAGTGTG |
| R MLV 7174 R | ACGTAACTGGGAGGATGGTAGGTG |
| R MLV 7845 R | CATTCCCCCCTTTTTCTGGAAACT |
| R MLV 8280 R | TGCAACAGCAAGAGGATTTATTG |
| KRM 595 | TTAATTGTCGACGCCACCATGGCGCGTTCAACGCTCTC |
| KRM 596 | TATATAGGCGCCGTTGGATCTCTCAAACAGGCCGTAAAC |
| KRM 597 | TTAATTGTCGACGCCACCATGGCGTGTTCAACGCTCTC |
| KRM 598 | TATATAGGCGCCATAGGATTTTTCAAACTGGCTATAGACG |
| KRM 605 | GTCCCGCTACTAGACTCAACCAGCTCTCTTGGGTTTTGTC |
| KRM 606 | GACAAAACCCAAGAGAGCTGGTTGAGTCTAGTAGCGGGAC |
| KRM 611 | GGTCCAGACACTAGGCTTAACCAGCTCTCTTGAGTTTTATCA |
| KRM 612 | TGATAAAACTCAAGAGAGCTGGTTAAGCCTAGTGTCTGGACC |

**S2 Table. List of antibodies and flow cytometry reagents used in this study.**

| Antibody | Clone | Dilution | Manufacturer | Catalog # |
|---|---|---|---|---|
| B220-PerCPCy5.5 | RA3-6B2 | 1:100 | BioLegend | 103236 |
| CD11b-APC | M1/70 | 1:300 | BD Biosciences | 553312 |
| CD11c-BV711 | HL3 | 1:300 | BD Biosciences | 563048 |
| CD138-BV786 | 281-2 | 1:100 | BD Biosciences | 569692 |
| CD19-BUV805 | 1D3 | 1:500 | BD Biosciences | 749027 |
| CD21/35-BV421 | 7G6 | 1:200 | BD Biosciences | 562756 |
| CD254-AF647 | IK22-5 | 1:100 | BD Biosciences | 560296 |
| CD279-BV786 | 29F.1A12 | 1:200 | BD Biosciences | 568566 |
| CD31-PerCP-Cy5.5 | 390 | 1:100 | BD Biosciences | 102419 |
| CD38-BUV737 | 90/CD38 | 1:100 | BD Biosciences | 741748 |
| CD3-APCCy7 | 17A2 | 1:150 | BioLegend | 100222 |
| CD3-PECy7 | 145-2C11 | 1:500 | BD Biosciences | 552774 |
| CD44-BV605 | IM7 | 1:300 | BD Biosciences | 563058 |
| CD45-PerCP | 30-F11 | 1:500 | BD Biosciences | 557235 |
| CD45-PETexasRed | 30-F11 | 1:400 | Invitrogen | MCD4517 |
| CD4-BUV563 | GK1.5 | 1:500 | BD Biosciences | 565709 |
| CD4-BUV737 | RM4-5 | 1:300 | BD Biosciences | 612844 |
| CD4-PECy7 | GK1.5 | 1:500 | ThermoFisher | 25-0041-82 |
| CD86-PE | PO3 | 1:100 | BioLegend | 105106 |
| CD8a-BUV395 | 53-6.7 | 1:100 | BD Biosciences | 553027 |
| CD8a-BUV570 | 53-6.7 | 1:500 | BioLegend | 100740 |
| CD95-PECy7 | JO2 | 1:100 | BD Biosciences | 557653 |
| CXCR4-BV421 | L276F12 | 1:200 | BioLegend | 146511 |
| CXCR5-AF647 | L138D7 | 1:100 | BioLegend | 145532 |
| FOXP3-eFluor450 | FJK-16s | 1:100 | ThermoFisher | 48-5773-80 |
| goat anti-mouse IgG (H+L) HRP | n/a | 1:5000 (ELISA) 1:10000 (Immunoblot) | Jackson ImmunoResearch | 115-035-146 |
| goat anti-mouse IgG1 HRP | n/a | 1:5000 | Jackson ImmunoResearch | 115-035-205 |
| goat anti-mouse IgG2a HRP | n/a | 1:5000 | Jackson ImmunoResearch | 115-035-206 |
| goat anti-mouse IgG2b HRP | n/a | 1:5000 | Jackson ImmunoResearch | 115-035-207 |
| goat anti-mouse IgG2c HRP | n/a | 1:2500 | Jackson ImmunoResearch | 115-035-208 |
| goat anti-mouse IgG3 HRP | n/a | 1:5000 | Jackson ImmunoResearch | 115-035-209 |
| goat anti-mouse IgM HRP | n/a | 1:5000 | Jackson ImmunoResearch | 115-035-075 |
| goat anti-rabbit IgG (H+L) HRP | n/a | 1:5000 (ELISA) 1:10000 (Immunoblot) | Jackson ImmunoResearch | 111-035-144 |
| IgG1-BUV395 | A85-1 | 1:300 | BD Biosciences | 740234 |
| IgG2a-AF594 | RMG2a-62 | 1:300 | BioLegend | 407119 |
| IgG2b-APC | RMG2b-1 | 1:300 | BioLegend | 406711 |
| IgM-FITC | II/41 | 1:300 | BD Biosciences | 553437 |

| | | | | |
|---|---|---|---|---|
| Ly-6G and Ly-6C-BV421 | RB6-8C5 | 1:500 | BD Biosciences | 562709 |
| MAdCAM-1-BUV563 | MECA-367 | 1:300 | BD Biosciences | 748769 |
| PDPN-PECy7 | 8.1.1 | 1:100 | BioLegend | 127411 |
| Tbet-PE | 4B10 | 1:100 | BD Biosciences | 561265 |
| TER-119-BUV395 | Ly-76 | 1:200 | BD Biosciences | 563827 |
| Zombie Aqua™ Fixable Viability Kit | n/a | 1:1000 | BioLegend | 423101 |
| Zombie NIR™ Fixable Viability Kit | n/a | 1:1000 | BioLegend | 423106 |
